# Generation of photocaged nanobodies for *in vivo* applications using genetic code expansion and computationally guided protein engineering

**DOI:** 10.1101/2021.04.16.440193

**Authors:** Jack M. O’Shea, Angeliki Goutou, Cyrus Sethna, Christopher W. Wood, Sebastian Greiss

**Affiliations:** Centre for Discovery Brain Sciences, University of Edinburgh, UK; Institute of Quantitative Biology, Biochemistry and Biotechnology, University of Edinburgh, UK

## Abstract

Nanobodies are becoming increasingly popular as tools for manipulating and visualising proteins *in vivo*. The ability to control nanobody/antigen interactions using light could provide precise spatiotemporal control over protein function. We develop a general approach to engineer photo-activatable nanobodies using photocaged amino acids that are introduced into the target binding interface by genetic code expansion. Guided by computational alanine scanning and molecular-dynamics simulations, we tune nanobody/target binding affinity to eliminate binding before uncaging. Upon photo-activation, binding is restored. We use this approach to generate improved photocaged variants of two anti-GFP nanobodies. These variants exhibit photo-activatable binding triggered by illumination with 365nm light. We demonstrate that the photocaged nanobodies we have created are highly robust and function in a complex cellular environment. We apply them to control subcellular protein localisation in the nematode worm *C. elegans*. Our approach provides a rare example of computationally designed proteins being directly applied in living animals and demonstrates the importance of accounting for *in vivo* effects on protein-protein interactions.

## Introduction

Nanobodies are small, 12-15kD, fragments of camelid heavy-chain only antibodies. Despite their small size, which corresponds to around 10% the size of standard antibodies, they exhibit potent and specific target binding. In recent years, nanobodies have been developed into indispensable tools for investigating protein function, in part due to the fact that they can be easily expressed *in vivo*.^[1,2]^ This unique set of properties has been leveraged to develop a variety of tools for labelling and manipulating proteins, to gain insights into biological processes. Fluorescently tagged nanobodies, termed “chromobodies”, have been used for *in vivo* antigen labelling.^[3]^ The ability of nanobodies to bind and interfere with specific protein domains makes them effective and highly specific inhibitors,^[4–6]^ capable of blocking pathogenic functions of proteins such as the *Clostridium difficile* toxin CDT,^[7]^ and the SARS-CoV-2 Spike.^[8,9]^ Besides direct inhibition, nanobodies have been used to interfere with antigen function by forced relocalisation of the antigen away from its subcellular region of activity,^[4]^ or by proteasomal degradation of the antigen via recruitment of the cellular ubiquitination machinery.^[10–14]^ The binding specificity of nanobodies and ease of intracellular expression has been exploited to improve ligand affinity to receptors for observation of rare binding events.^[15]^ Additionally, nanobodies have been used to deliver photo-switchable ligands to receptors for optical control of receptor activation.^[16]^ Their small recognition epitopes even facilitate tools wherein two nanobodies that bind different regions of GFP can control transcription by using GFP as a bridge to bring together split proteins, such as split transcription factors and split Cre recombinase.^[17,18]^ Nanobodies have been employed as tools in a wide range of model systems, including mammalian cells, mice, zebrafish, and *C. elegans*.^[2]^

A limitation of native nanobodies is that their binding cannot be modulated after expression. The ability to induce antigen binding would significantly increase the applicability of nanobody-based tools by adding a level of spatiotemporal control absent when using native nanobodies. Several avenues have been explored to install the ability to control nanobody/antigen interaction. Insertion of a ligand-binding domain into the loop regions of various nanobodies has been used to impart chemo-inducibility and make antigen binding dependent on small molecule ligands.^[19]^ However, chemical control of nanobody binding does not offer spatial control and offers only limited temporal control due to slow diffusion kinetics of the required chemical ligand, especially in multicellular systems. Optical control is an attractive alternative to chemical control, since it potentially allows high-precision spatiotemporal manipulation.

Three avenues to install optical control in nanobodies have been explored. The fusion of split nanobody fragments with photo-inducible dimerisers allows photoinducible binding, however this approach can be limited by slow kinetics, and binding saturation is only achieved after extended illumination.^[20]^ Insertion of photo-switchable domains into nanobody loops also achieved photoinducible antigen binding or dissociation, but the direction and magnitude of the effect can be unpredictable.^[21]^ These two approaches also suffer from requiring blue light for induction, which can interfere with imaging and also makes them incompatible with other blue light dependent optogenetic tools.^[14,16,22]^ The third approach involves the use of genetic code expansion^[23]^ to incorporate photocaged amino acids (Figure 1A) into the binding interface of the nanobody with its antigen, thereby preventing binding until the caging group is removed by illumination (Figure 1B, 1C). Uncaging of photocaged amino acids is rapid, can be performed with excellent spatial resolution, and the wavelengths involved are compatible with most imaging and optogenetics techniques.^[24]^

**Figure 1.**
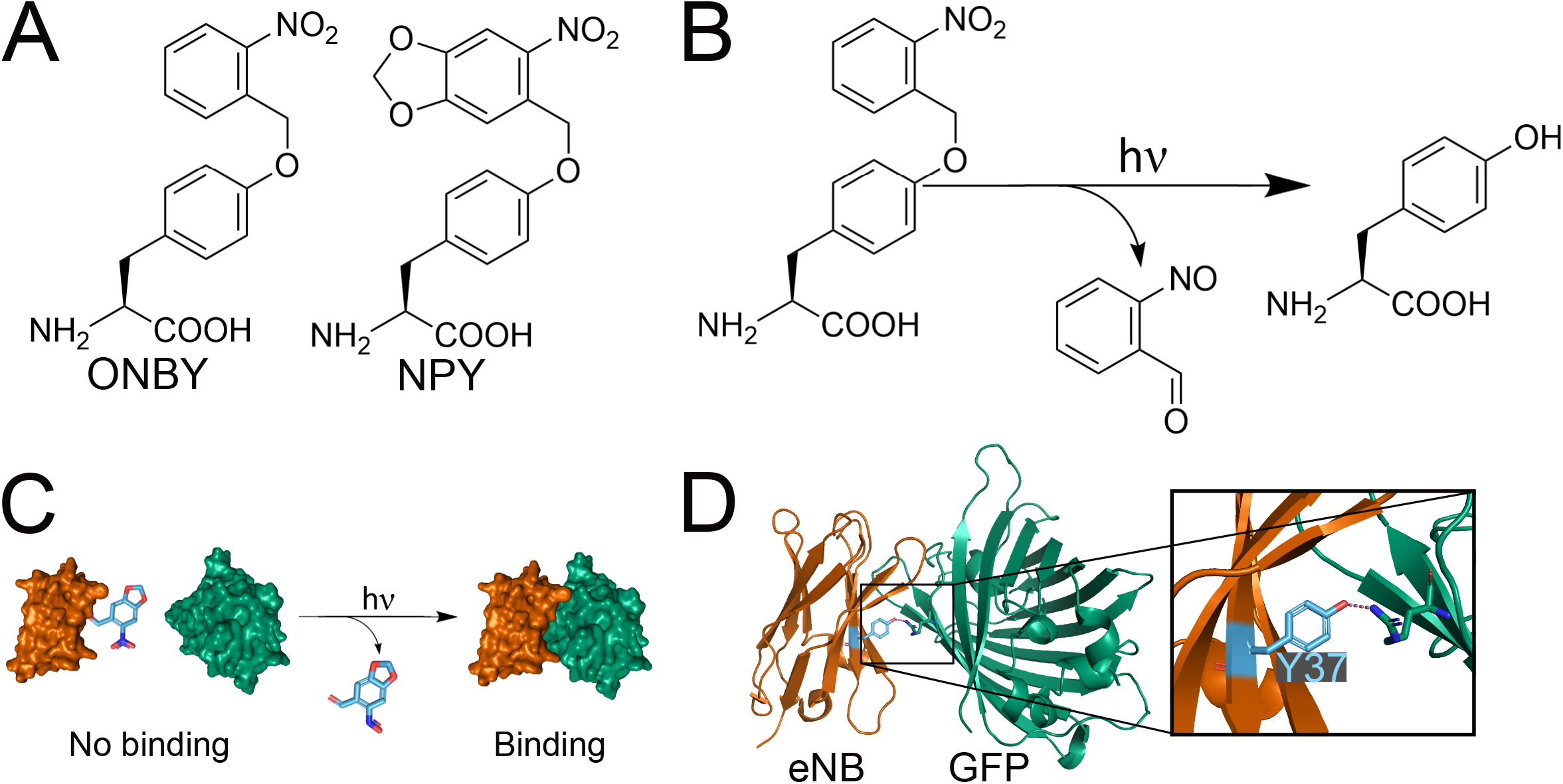
**A**. Chemical structures of the photocaged tyrosines ortho-nitrobenzyl tyrosine (ONBY) and nitropiperonyl tyosine (NPY). **B**. The uncaging reaction of ONBY. Uncaging of NPY follows the same mechanism. **C**. Schematic for photoinducible protein-protein interactions using photocaged amino acids. The photocaging group (in blue) prevents interaction of the binding partners, but is removed by illumination. **D**. Crystal structure (PDB: 3K1K) of the interaction of the anti-GFP “enhancer” nanobody (eNB) and GFP. Highlighted in blue is Y37 of eNB, which forms a polar interaction with R168 of GFP.

Substituting key antigen binding residues on nanobodies with photocaged amino acids has been shown to reduce nanobody/antigen binding *in vitro*.^[25,26]^ However, it is unknown whether these reductions are sufficient to abolish binding *in vivo* when both the photocaged nanobody and its antigen are expressed *in situ*. To date, the closest attempt was the delivery of purified photocaged nanobodies synthesised in *E. coli* to HeLa cells expressing the antigen.^[27]^ Factors influencing nanobody/antigen interaction such as pH, ionic concentration, incubation time, and intracellular concentration of nanobody and antigen may differ significantly from *in vitro* and delivered conditions.^[28]^ The disruption of binding by the photocaged amino acid observed in these cases may therefore not correctly predict binding when directly expressing photocaged nanobodies *in vivo*. The difference between intracellular and *in vitro* conditions could result in failure of the photocaged amino acid to abolish the nanobody/antigen interaction *in vivo*. In this case, a solution would be the modification of the binding interface through the introduction of further mutations to tune the binding strength for the desired environment.

Computational Alanine Scanning (CAS) is a generally applicable technique for identifying hot-spot residues or sets of residues that contribute to binding interactions. CAS programs are *in silico* alternatives to classic alanine scanning mutagenesis experiments, which can be prohibitively time-consuming and costly. BUDE Alanine Scan (BudeAlaScan) is an open-source, state-of-the-art CAS program freely available for use via the in-browser application BAlaS.^[29,30]^ BudeAlaScan calculates the binding free energy of a given protein-protein interaction (PPI), for the native protein sequence and for when each residue is independently mutated to alanine. The software accepts protein crystal structures or multi-model files such as NMR ensembles and molecular-dynamics trajectories.

Here we describe the rational engineering of nanobody paratopes to tune *in vivo* binding to their target protein. This allows us to optically control binding through incorporation of the photocaged tyrosine variants ONBY and NPY (Figure 1A). We show that the introduction of a photocaged non-canonical amino acid (ncAA) alone is not sufficient to break nanobody/antigen binding *in vivo*. We perform CAS to identify surface residues on the nanobodies that contribute to binding and use this information to guide engineering of nanobodies by introducing alanine mutations in place of interaction hot-spot residues so that binding is abolished in the photocaged form and restored on uncaging. We demonstrate the general applicability of our approach by engineering two anti-GFP nanobodies and using them to control protein localisation in a living animal, the nematode worm *C. elegans*.

## Results

### Incorporating photocaged tyrosine in *C. elegans*

We set out to photocage nanobodies for use in *C. elegans*, initially focussing on a well-established anti-GFP nanobody, the “enhancer” nanobody (eNB, Figure 1D),^[31]^ as it is utilised for many *in vivo* tools^[11,12,16–18]^ and has successfully been expressed and shown to bind GFP in *C. elegans*.^[11]^ Ortho-nitrobenzyl tyrosine (ONBY) (Figure 1A), a photocaged form of tyrosine, has been substituted for Y37, located in the GFP binding interface of eNB (Figure 1D), and was shown by a cell-surface binding assay to reduce binding affinity by 10,000 fold.^[25]^ In contrast, the alternative photocaged tyrosine nitropiperonyl tyrosine (NPY) (Figure 1A) was previously found to only reduce binding by 420 fold.^[25]^ The nitropiperonyl group on NPY has a red shifted absorption maximum compared to the ortho nitro group on ONBY, making it preferable for *in vivo* applications due to the resulting improved uncaging and thus shorter required illumination times at 365nm.^[32]^ Neither ONBY nor NPY has been used to control a nanobody binding *in vivo* when expressed *in situ* with its antigen.

To express photocaged nanobodies *in vivo*, we first established a system for incorporation of photocaged tyrosine in *C. elegans*. Genetic code expansion utilises orthogonal aminoacyl synthetase / tRNA pairs to site specifically direct incorporation of non-canonical amino acids (ncAA) into target proteins. The most widely applied orthogonal genetic code expansion system is based on the pyrrolysyl synthetase (PylRS) / tRNA(Pyl) from methanogenic archaea, which has been used to expand the genetic code of bacteria, eukaryotic cultured cells, plants, and animals.^[33]^ In *C. elegans*, incorporation of ncAA is possible in all tissues and photocaged amino acids have been used to control protein function in living animals.^[34,35]^

Several caging groups have been used to photocage tyrosine and orthogonal aminoacyl-tRNA-synthetase / tRNA pairs have been described in the literature to incorporate them,^[32,36]^ albeit thus far not in animals. We decided to use the pyrrolysyl-tRNA synthetase variant NBYRS, which can be used to incorporate both ONBY and NPY amino acids.^[32]^ We introduced the previously described mutations into a Methanosarcina mazei (Mm) PylRS gene optimised for expression in *C. elegans* to generate MmNBYRS.

To improve MmNBYRS tRNA acylation efficiency, we first added a nuclear-export sequence (NES) derived from human PKIα to the N-terminus of MmNBYRS to increase cytoplasmic localisation.^[35,37]^ Second, in place of wild-type tRNA(Pyl)_CUA_ we used the tRNA(C15)_CUA_ variant, which improves incorporation efficiency in mammalian cells and in *C. elegans*.^[35,38]^ We showed previously that, in combination, these two improvements can increase incorporation of ncAA in *C. elegans* by more than 50-fold.^[35]^

We generated transgenic strains ubiquitously expressing a fluorescent reporter for incorporation along with the genetic code expansion machinery components NES-PKIα::MmNBYRS and tRNA(C15)_CUA_. The reporter consists of an N-terminal GFP separated by a TAG codon from a C-terminal mCherry fused to the *C. elegans* EGL-13 nuclear localisation sequence (NLS)^[39]^ followed by an HA tag. The reporter is designed so that prior to incorporation, only GFP is expressed, which is localised throughout the cell, while incorporation of a ncAA at the TAG codon will result in the production of full length GFP::mCherry::NLS protein that is localised to the nucleus (Figure 2A).

**Figure 2.**
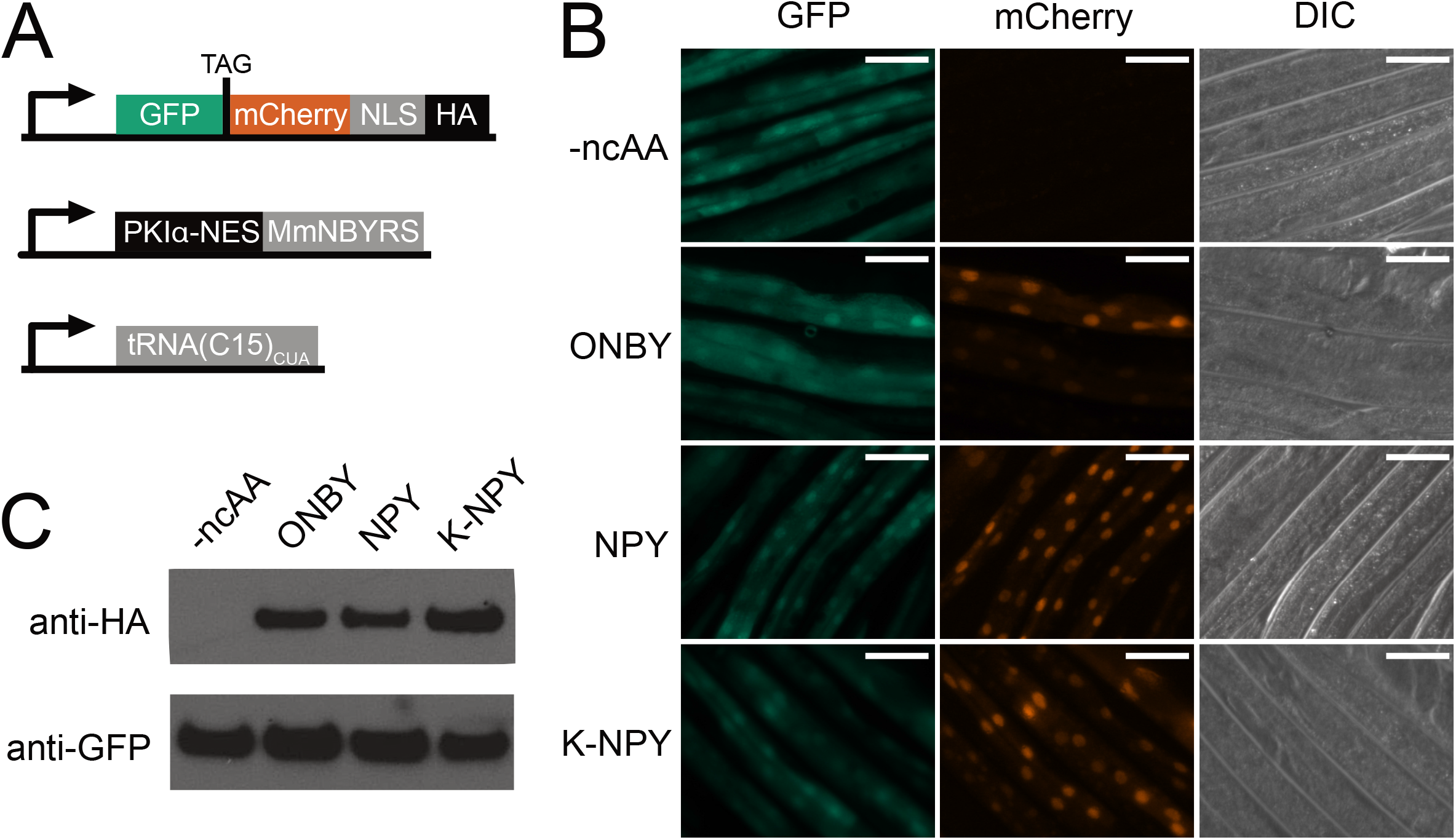
**A**. Genetic constructs for the fluorescent ncAA incorporation reporter and the machinery for incorporation of photocaged tyrosines. Successful incorporation at the TAG codon will result in the production of full length GFP::mCherry fusion protein. **B**. Fluorescence imaging of *C. elegans* expressing the fluorescent reporter construct and the ncAA incorporation machinery, in the absence of ncAA and in the presence of 0.1mM ONBY, NPY, or K-NPY. Scale bars 50µm. **C**. Anti-GFP and anti-HA Western blots performed on lysates of the *C. elegans* shown in (B), grown in the absence of ncAA or in the presence of 0.1mM ONBY, NPY, or K-NPY.

Since the reporter construct contains an internal stop codon, it is a target for nonsense mediated decay (NMD). In order to increase the amount of reporter mRNA and ensure easy visualization of incorporation, we generated transgenic strains in the *smg-6(ok1794)* deletion background, which makes the animal deficient for NMD.

We tested incorporation efficiency by transferring transgenic animals to Nematode Growth Medium (NGM) agar plates containing ncAA (0.1 mM ONBY or 0.1mM NPY). We observed strong red fluorescence in the nucleus, appearing within 24 hours after introduction to ncAA, indicating successful incorporation (Figure 2B). We further confirmed the identity of the full-length reporter protein by western blot using antibodies against the C-terminal HA tag (Figure 2C, Supplementary Figure 1).

We found that both the ONBY and NPY amino acids were incorporated with comparable efficiency (Figure 2C). Previous reports indicate that supplying ncAA to *C. elegans* as dipeptides may increase uptake of the compounds.^[40]^ We therefore supplied NPY as a K-NPY dipeptide with lysine attached to the NPY amino group. However, we did not see increased incorporation for K-NPY as compared to NPY (Figure 2C). Nevertheless, we found that in our buffer conditions NPY was not soluble at concentration above 0.1mM, while K-NPY was easily soluble into the mM range. While the dipeptide did not improve amino acid incorporation efficiency, higher ncAA concentrations may help to increase production of photocaged proteins in cases where baseline incorporation efficiency is low. In all following experiments NPY is supplied to the animals in the dipeptide form K-NPY.

### NPY and ONBY substitution is not sufficient to photocage antigen binding *in vivo*

We proceeded to express the anti-GFP nanobody eNB containing either of the photocaged tyrosines ONBY or NPY in lieu of the native Y37 residue, which is situated at the binding interface.^[25]^ Since both ncAA showed efficient incorporation, we decided to construct transgenic strains for eNB expression in the wild type N2 strain, which has functional NMD.

To assay eNB to GFP binding *in vivo*, under physiological conditions, we employed a subcellular localisation assay as a visual read-out. We expressed a eNB::mCherry fusion protein with mCherry attached to the eNB C-terminus. We furthermore co-expressed GFP fused to the *C. elegans* EGL-13 nuclear localisation signal (NLS)^[39]^ (Figure 3A), so that binding of eNB::mCherry to GFP would result in the nuclear localisation of the red mCherry signal. We reasoned that GFP binding strength of different eNB variants (with and without photocaged residues) could be compared by measuring the nuclear to cytoplasmic ratio (N/C) of the eNB::mCherry fusion. Strong binders would be mostly nuclear, whereas non-binders would be free to diffuse throughout the cell (Figure 3B). The eNB::mCherry fusion has a molecular weight of around 40kDa, a size which easily allows free diffusion across the nuclear membrane.^[41]^ Additionally, the C-terminal mCherry would act as a reporter for successful ncAA incorporation. We confirmed that the wild-type eNB::mCherry (i.e. not containing any photocaged residues) indeed localises to the nucleus *in vivo* and therefore binds to GFP::NLS. Conversely, co-expression of eNB::mCherry with NLS::mTagBFP2, a blue fluorescent protein which is not related to GFP, does not lead to nuclear localisation of eNB::mCherry. Likewise, mCherry alone, without a fused eNB, does not localise to the nucleus when co-expressed with GFP::NLS (Supplementary Figure 2).

**Figure 3.**
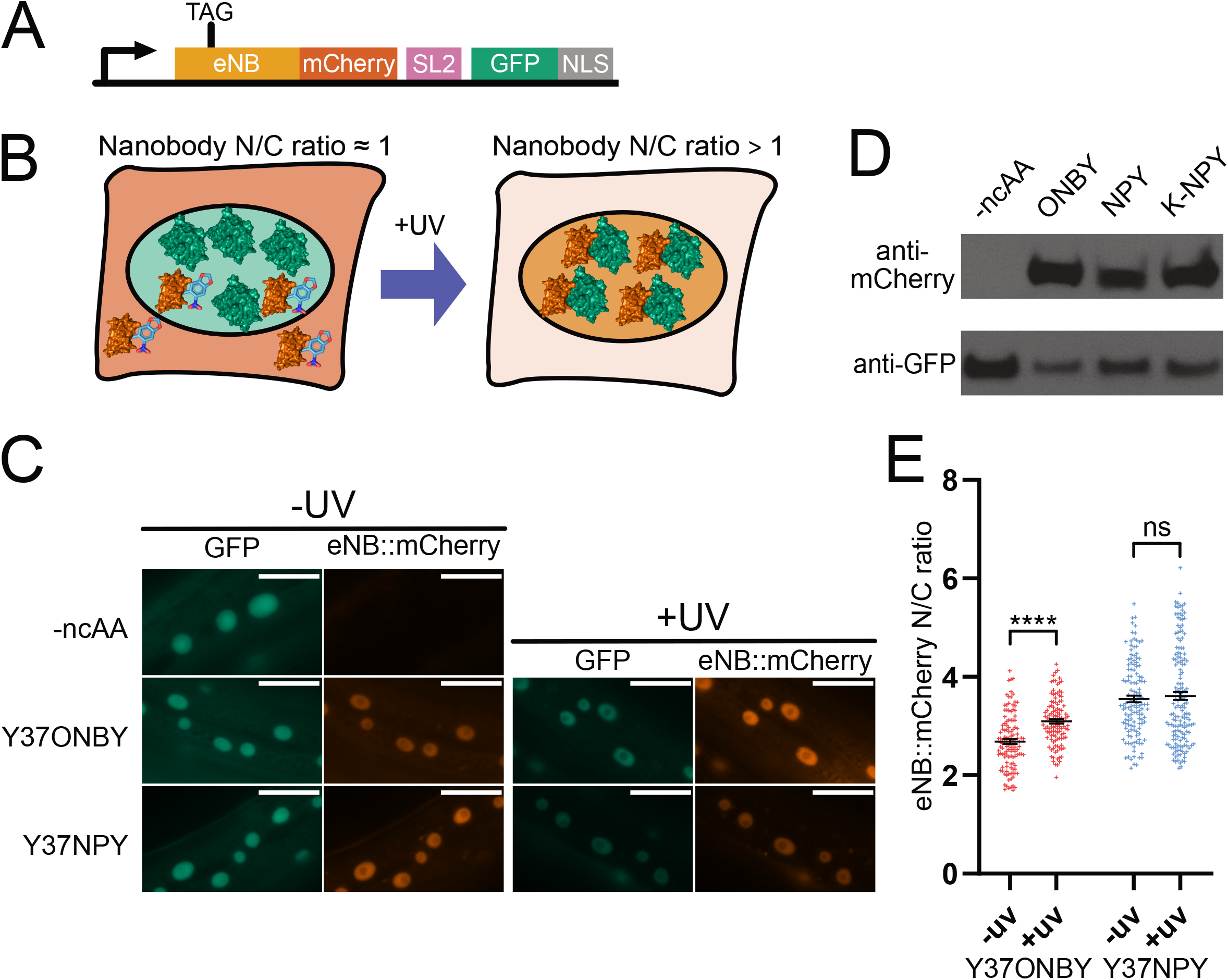
**A**. Genetic construct for the eNB/GFP interaction reporter. eNB::mcherry is produced upon incorporation of ncAA at the TAG codon. GFP fused to a nuclear localisation sequence is expressed in an artificial operon behind the same promoter independently of ncAA incorporation. **B**. Schematic of the *in vivo* photoinducible interaction assay. Photocaged nanobodies fused to mCherry are co-expressed with nuclear GFP. If incorporation of a photocaged amino acid into the GFP-binding region of the nanobody is sufficient to abolish GFP binding, the mCherry nuclear/ cytoplasmic ratio (N/C) will be near 1, as measured by mCherry fluorescence. Following uncaging with 365nm light (+UV), the nanobody will regain its ability to bind GFP and relocate to the nucleus, leading to and increase in the N/C ratio. **C**. Images of cells in animals of the *in vivo* assays of the interaction between GFP and eNB^wt^, eNB^Y37ONBY^, or eNBY37NPY, before and after 365nm illumination. Scale bars 20µm. **D**. Anti-GFP and anti-mCherry Western blots performed on lysates from *C. elegans* expressing the interaction reporter constructs described in 3A and grown in the absence of ncAA or in the presence of 0.1mM ONBY, NPY, or K-NPY. **E**. Quantification of mCherry nuclear/ cytoplasmic ratio for eNB with mutations Y37ONBY or Y37NPY. Data are presented as measurements of individual cells and mean ± SEM. Measurements were taken from 7-10 animals per condition. ns p > 0.05; **** p < 0.0001.

To direct incorporation of ONBY and NPY, we introduced a TAG stop codon replacing the codon for Y37 into eNB. We co-expressed eNB(TAG)::mCherry together with GFP::NLS and the components of the photocaged tyrosine incorporation machinery, consisting of NES-PKIα::MmNBYRS and tRNA(C15)_CUA_.

The transgenic *C. elegans* strains we generated showed a strong GFP signal that was exclusively nuclear. As expected, we saw no mCherry signal in the absence of ncAA. When we transferred the animals to plates supplemented with 0.1 mM ONBY, NPY, or K-NPY we saw clearly visible red fluorescence appearing within 24h, indicating production of the full length eNB^Y37ONBY^::mCherry or eNB^Y37NPY^::mCherry fusion proteins, we confirmed the production of full length protein by western blot for mCherry (Figure 3C, 3D, Supplementary Figure 1). As with the GFP::mCherry incorporation reporter we saw no difference in incorporation efficiency between the ncAAs, even when supplying NPY as the dipeptide K-NPY.

To our surprise, we found that both photocaged nanobodies eNB^Y37ONBY^ and eNB^Y37NPY^ co-localised with GFP::NLS in the nucleus (Figure 3C). This observation was very unexpected since previously published *in vitro* binding assays, using photocaged eNB purified from *E. coli*, showed that Y37 substitution for ONBY reduces binding affinity 10,000 fold and blocks binding *in vitro*, and NPY reduces binding affinity 420 fold.^[25]^ However, while ONBY was not sufficient to abolish binding in our intracellular assay, we found that uncaging by illuminating animals using a 365nm LED (640 seconds illumination, 10mW/cm^2^) led to a clear increase in the N/C ratio for eNB^Y37ONBY^ from 2.69 (± 0.05) to 3.10 ± (0.04) (Figure 3E, Supplementary Figure 3). This contrasts with eNB^Y37NPY^, where there was no significant change in N/C, going from 3.55 (± 0.07) before illumination to 3.61 (± 0.08) after uncaging. Our *in vivo* observations therefore support the previous *in vitro* observation that ONBY is more disruptive to the interaction than NPY. The disruption by ONBY is however not sufficient to block binding *in vivo*.

### Molecular-dynamics simulations and *in silico* alanine scan for tuning of eNB/GFP interaction

We reasoned that in order to generate photoinducible nanobodies for *in vivo* use, it would be necessary to further engineer the nanobody/antigen binding interface. Our aim was to reduce the binding strength so that the introduction of a photocaged amino acid would abolish binding, yet upon uncaging the nanobody would retain sufficient affinity to bind its target.

Since nanobodies bind their antigen via multiple interactions, we hypothesised that removing interactions between eNB and GFP in addition to Y37 photocaging would result in an eNB variant optimised for *in vivo* photoinducible GFP binding (Figure 4A). To develop a universally applicable approach to identify suitable candidate residues to tune the binding strength, we turned to computational alanine scanning (CAS).

**Figure 4.**
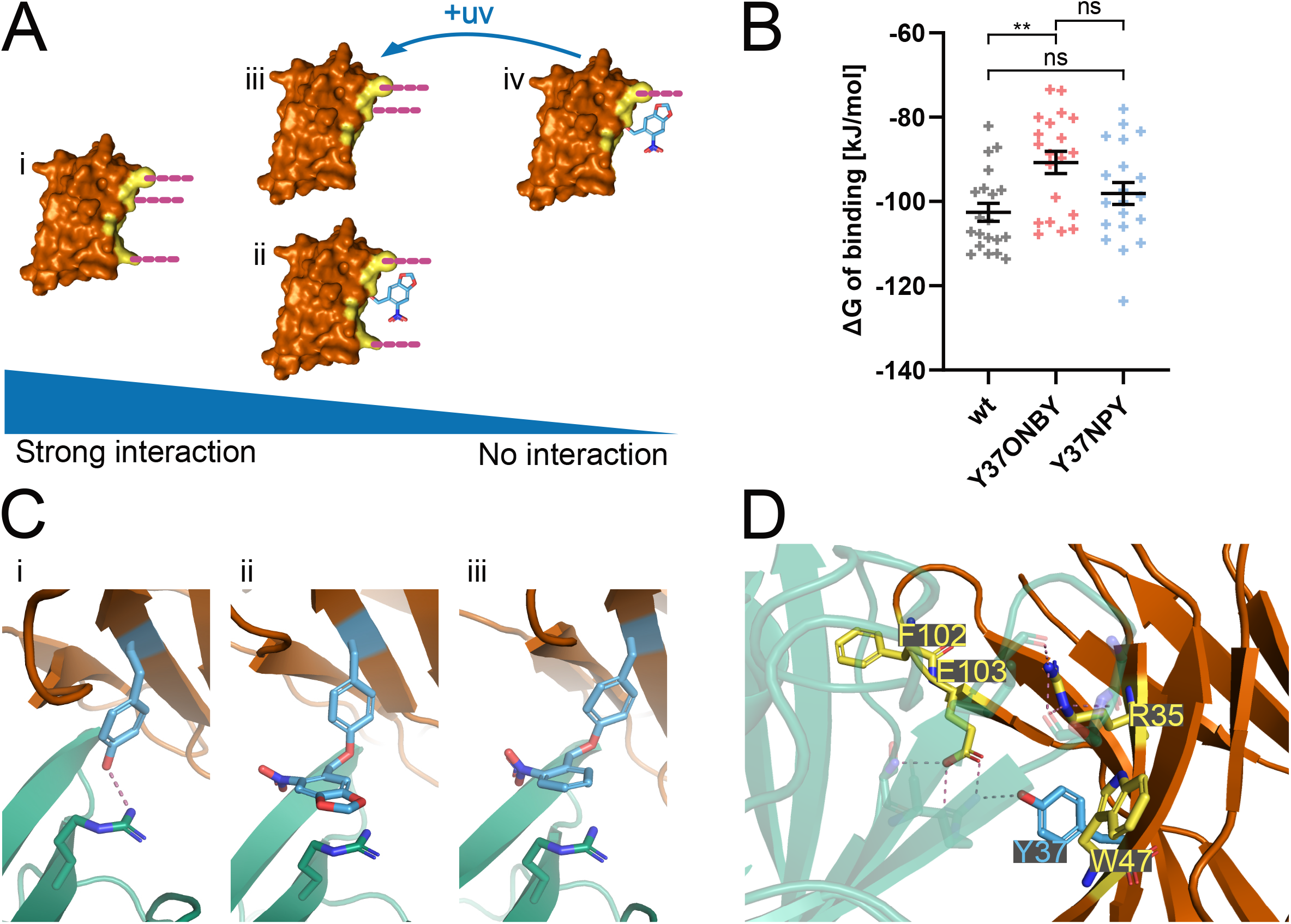
**A**. Schematic for the design of an improved photocaged nanobody. The native nanobody **(i)** makes several interactions with its antigen. Photocaging **(ii)** or mutating **(iii)** one of these interactions is insufficient to abolish the interaction. Introducing a photocaging group and a mutation to the nanobody abolishes the interaction **(iv)**. The interaction can be restored by removing the caging group through illumination with 365nm. **B**. BUDE Alanine Scan (BAlaS) calculations of ΔG of binding for the interaction between GFP and eNB^wt^, eNB^Y37ONBY^, or eNB^Y37NPY^. Data are presented as individual simulations and mean ± SEM. ns p > 0.05; ** p < 0.01. **C**. Representative frames from MD trajectories for the interaction between R168 of GFP and Y37 **(i)**, NPY37**(ii)**, or ONBY37**(iii)** of eNB. **D**. eNB/GFP interface (PDB: 3K1K) with eNB residues that are predicted by BAlaS to have the greatest effect on interaction strength when mutated to alanine are highlighted in yellow.

Bude Alanine Scan (BAlaS) is a CAS program that calculates the binding free energy (ΔG) of a native PPI, and the change in binding free energy that occurs when residues are independently mutated to alanine. We used BAlaS to perform *in silico* alanine scanning on eNB in complex with GFP to generate a shortlist of alanine mutations predicted to decrease binding strength. We hypothesised that introducing such alanine mutations into eNB would reduce binding affinity and this, in combination with photocaging of Y37, would make the eNB/GFP interaction photoinducible.

BAlaS has been demonstrated to generate more accurate predictions of hot-spot residues when multiple conformations of the protein are provided.^[30]^ To generate input data for BAlaS, we first ran twenty independent 20ns molecular-dynamics (MD) simulations for each nanobody variant: eNB^wt^, eNB^Y37ONBY^, and eNB^Y37NPY^. Electronic parameters for the ONBY and NPY residues were derived using the Generalised Amber Force Field,^[42]^ except for the nitro group, where values from quantum mechanical calculations were used.^[43]^ Simulations were run using the OpenMM software.^[44]^ The simulations rapidly reach equilibrium, after which point the temperature and energy remains constant (Supplementary Figure 4). The final frames from each of the twenty simulations were used as inputs for BAlaS, representing a sample of twenty representative poses the complex can adopt. The output ΔG value for each variant is the mean binding free energy of these twenty poses.

BAlaS predicted ΔGs of the three eNB variants eNB^wt^, eNB^Y37ONBY^, and eNB^Y37NPY^ that are in line with the experimental results of ourselves (Figure 4B) and others.^[25]^ As expected, eNB^wt^ is predicted to have the strongest interaction with GFP, with a mean ΔG of −102.60kJ/mol (± 2.07). Of the photocaged variants, the mean ΔG predictions are −90.73kJ/mol (± 2.54) for eNB^Y37ONBY^ and −98.09kJ/mol (± 2.54) for eNB^Y37NPY^. Interestingly, the ΔΔG of 11.87 kJ/mol (p=0.0012) between eNB^wt^ and eNB^Y37ONBY^ is more pronounced as compared to the smaller ΔΔG of 4.51kJ/mol (p=0.19) between eNB^wt^ and eNB^Y37NPY^. This is in agreement with the findings that ONBY is more disruptive to the interaction than NPY, observed both *in vitro* by Jedlitzke et al.^[25]^ and *in vivo* by ourselves, supporting the robustness of our *in silico* predictions.

Close examination of the MD trajectories offers explanations for why neither ONBY nor NPY abolish binding, and why ONBY is more disruptive than the bulkier NPY (Figure 3E). In the crystal structure and MD trajectories of the eNB^wt^/GFP interaction, Y37 of eNB forms a hydrogen bond with R168 of GFP (Figure 4C, panel i). This makes Y37 an obvious candidate for photocaging, and in every eNB^Y37ONBY^ and eNB^Y37NPY^ trajectory this hydrogen bond is abolished. However, in 38 out of the 40 photocaged eNB trajectories the photocaging groups form a stable cation-pi interaction with the positively charged guanidinium group of R168 on GFP, the two groups aligning in parallel to one another and packing tightly (Figure 4C, panels ii, iii). This interaction compensates for the loss of the Y37/R168 hydrogen bond, which may account for the poor disruptive effect of these caging groups.

To identify further eNB residues important for GFP binding, which could be combined with caged Y37 to make the interaction photoinducible, we performed a computational alanine scan on eNB^wt^ using BAlaS. The scan identified 28 eNB residues that are predicted to lower binding strength when mutated to alanine. The effects on the binding energy ranged from +0.01kJ/mol to +9.62kJ/mol (Supplementary Figure 5). The most disruptive alanine mutations predicted were R35A, Y37A, W47A, F102A and E103A, all of which are situated at the eNB/GFP interface (Figure 4D).

### Use of engineered photocaged eNB variants *in vivo*

To test whether the combination of photocaging and interface modification through the introduction of targeted alanine mutations could indeed be used to engineer *in vivo* photocaged nanobodies, we combined the disruptive alanine mutations R35A, W47A, or E103A, with a TAG codon at position Y37 to direct incorporation of photocaged tyrosines. We also generated a triple mutant combining the R35A and E103A mutations with the TAG mutation at Y37. We then constructed transgenic *C. elegans* strains expressing the MmNBYRS / tRNA (C15)_CUA_ ncAA incorporation machinery together with the eNB::mCherry mutants and GFP::NLS.

We decided to test eNB variants using the NPY caged tyrosine, since it is more amenable to uncaging at 365 nm than ONBY,^[32]^ and would therefore be preferable for *in vivo* usage. To incorporate NPY, we transferred animals to plates supplemented with 0.1mM K-NPY. 24h after transfer, we observed the appearance of red fluorescence indicative of NPY incorporation and production of full length eNB::mCherry protein.

In stark contrast to animals expressing eNB^Y37ONBY^ and eNB^Y37NPY^, where the red eNB::mCherry fluorescence was predominantly localised to the nucleus, we found that for the variants eNB^Y37NPY, R35A^, eNB^Y37NPY, E103A^, and eNB^Y37NPY, R35A, E103A^ the red fluorescence was distributed throughout the cell, indicating that the introduction of additional mutations were sufficient to abolish eNB to GFP binding. Interestingly, the eNB^Y37NPY, W47A^ variant retained strong GFP binding as evidenced by its clear nuclear localisation (Figure 5A).

**Figure 5.**
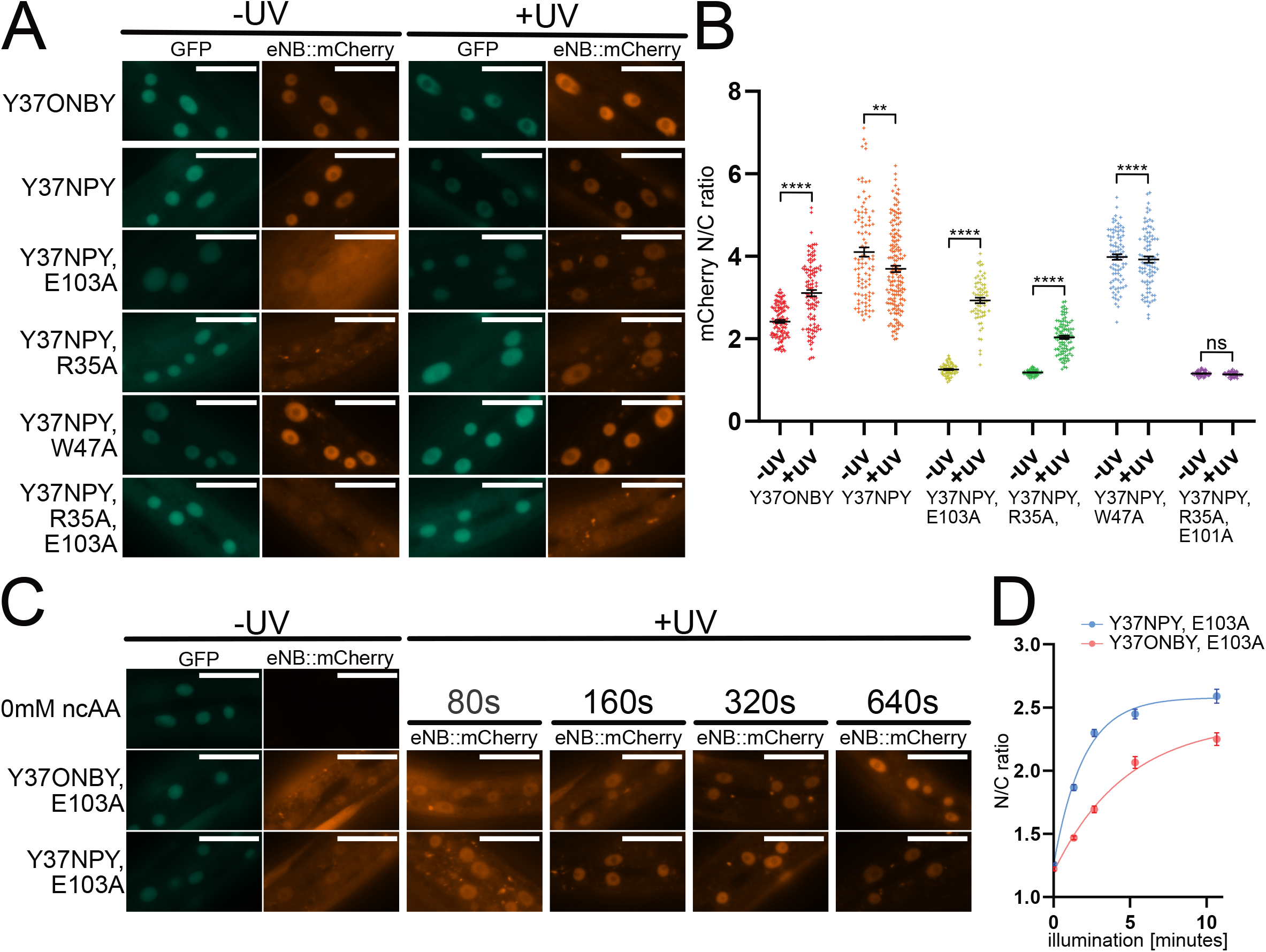
**A**. Images of cells in animals expressing different photocaged eNB variants, before and after 365nm illumination. Scale bar 20µm. **B**. Quantification of mCherry nuclear/cytoplasmic ratio of photocaged eNB variants before and after 365nm illumination. Data are presented as individual cell measurements and mean ± SEM. Measurements were taken from 7-10 animals per condition. ns p > 0.05; ** p < 0.01; **** p < 0.0001. **C**. Images of cells in animals expressing eNB^Y37ONBY, E103A^ and eNB^Y37NPY, E103A^, subjected toa range of 365nm illumination times. Scale bar 20µm. **D**. Quantification of mCherry nuclear/cytoplasmic ratio of eNBY37ONBY, E103A and eNBY37NPY, E103A subjected to a range of 365nm illumination times. Data are presented as mean ± SEM. Measurements were taken from 7-10 animals per condition. ONBY vs NPY: pre-UV p= 0.2151ns, all other time points p<0.0001; ONBY 640s vs NPY 160s p = 0.4110.

We proceeded to test whether removal of the caging group would re-establish eNB/GFP binding by quantifying the eNB::mCherry N/C ratio before and after uncaging. We illuminated animals using a 365 nm LED and compared these with unilluminated animals. Upon illumination we observed a striking shift in localisation to the nucleus for the eNB^Y37NPY, R35A^ and eNB^Y37NPY, E103A^ variants (Figure 5A, 5B). Illumination induced an increase in the N/C ratio of eNB^Y37NPY, R35A^ from 1.18 (± 0.01) to 2.04 ± (0.04), and of eNB^Y37NPY, E103A^ from 1.26 (± 0.02) to 2.59 (± 0.07) (Figure 5B, Supplementary Figure 6). eNB^Y37NPY, W47A^ remained nuclear with no significant change, going from 3.98 (± 0.07) to 3.92 (± 0.07). eNB^Y37NPY, R35A, E103A^ remained cytoplasmic following uncaging, its N/C ratio going from 1.16 (± 0.01) to 1.14 (± 0.01), indicating that introduction of two alanine mutations abolished the ability of the eNB variant to bind GFP even after removal of the photocaging group.

We then compared ONBY and NPY to test the effect of different photocaging groups on the effectiveness of our approach of tuning the interaction strength. For this, we grew animals expressing the eNB^Y37TAG, E103A^ variant on NGM agar plates supplemented with either 0.1mM ONBY or 0.1mM K-NPY. In both cases we observed the appearance of red fluorescence that was distributed throughout the cell, indicating the presence of eNB^Y37ONBY, E103A^ or eNB^Y37NPY, E103A^ (Figure 5C). Interestingly, there was no significant difference between N/C ratios of eNB^Y37ONBY, E103A^ and eNB^Y37NPY, E103A^ before uncaging, with values of 1.23 (± 0.01) and 1.26 (± 0.01) respectively (Figure 5D, Supplementary Figure 7). Upon uncaging using a 365 nm LED, red fluorescence accumulated in the nucleus for both variants. However, we saw a pronounced difference in the uncaging kinetics, with ONBY requiring significantly longer illumination times than NPY. eNB^Y37ONBY, E103A^ required 640s illumination to reach the N/C ratio of 2.26 (± 0.05), while eNB^Y37NPY, E103A^ reached an N/C ratio of 2.30 (± 0.03) with a quarter of the illumination time. This reflects the fact that NPY has a red-shifted absorption maximum compared to ONBY and is therefore more amenable to uncaging using 365nm.^[32]^

### Development of photocaged mNB

After successfully modifying eNB, we proceeded to test whether our approach was generally applicable by engineering a second nanobody, the “minimiser” GFP-binding nanobody (mNB).^[31]^ We chose Y116 on mNB as a candidate residue for substitution with a photocaged tyrosine to achieve photoinducible GFP binding. Y116 is in the centre of the binding interface and forms hydrogen bonds with GFP N164 and mNB Q102 (Figure 6A). The introduction of ONBY in place of this residue has previously been shown by Jedlitzke et al to moderately affect binding *in vitro*.^[25]^

**Figure 6.**
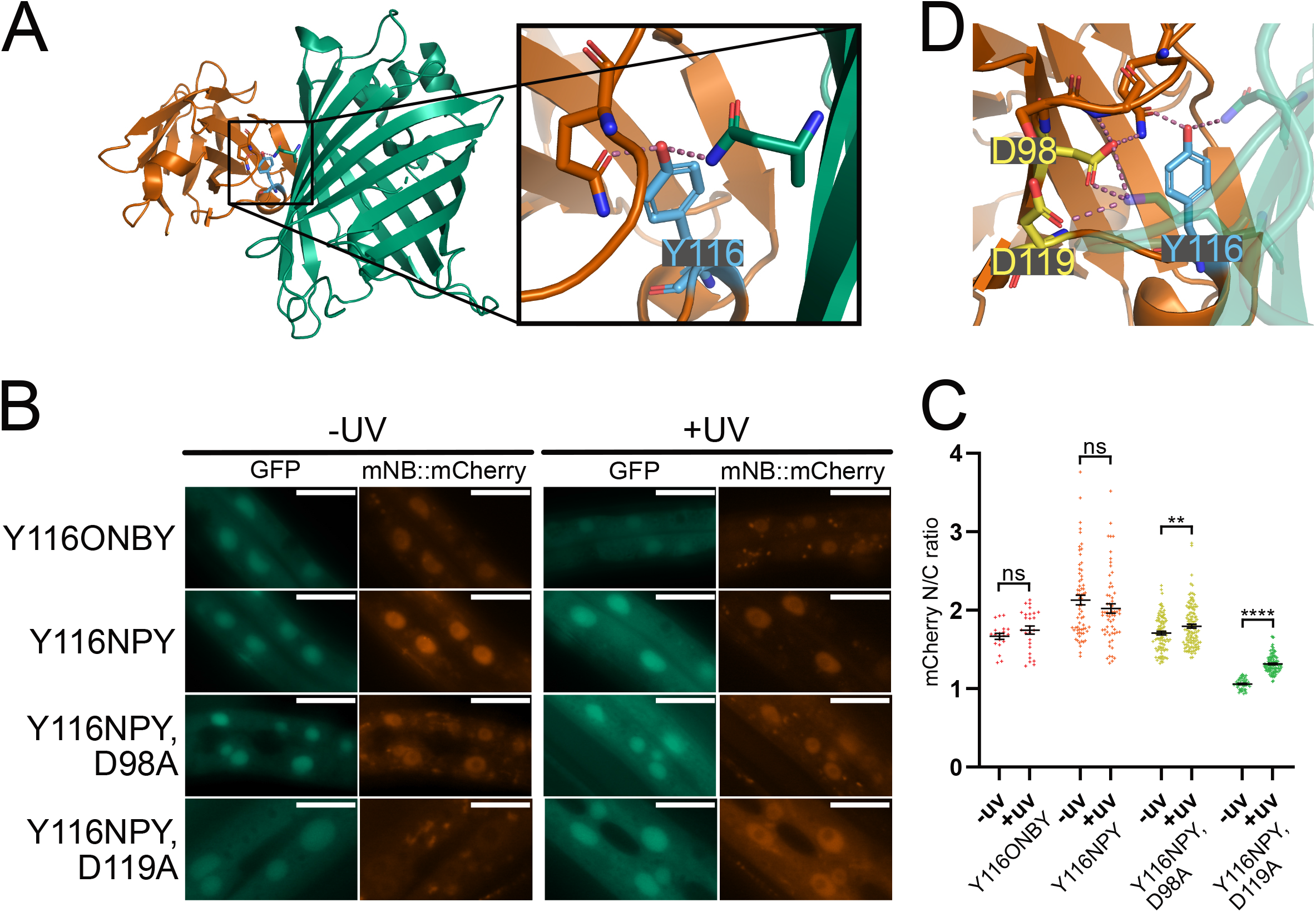
**A**. Crystal structure of mNB/GFP interaction (PDB: 3G9A). Highlighted in blue is Y116 of mNB, which forms a polar interaction with R168 of GFP. **B**. Localisation of photocaged mNB variants before and after 365nm illumination. Scale bar of 20µm. **C**. Quantification of mCherry nuclear/cytoplasmic ratio of photocaged mNB variants before and after 365nm illumination. Data are presented as individual cell measurements and mean ± SEM. Measurements were taken from 7-10 animals per condition. ns p > 0.05; ** p < 0.01; **** p < 0.0001. **D**. mNB/GFP interface (PDB: 3G9A) with two mNB residues that are predicted by BAlaS to have significant effect on interaction strength when mutated to alanine highlighted in yellow.

We specified incorporation of photocaged amino acids by introducing a TAG codon at position Y116 and generated *C. elegans* strains co-expressing the mNB^Y116TAG^ variant together with the photocaged tyrosine incorporation machinery. To quantify GFP binding of the photocaged mNB we again used our *in vivo* assay where photocaged mNB::mCherry fusion is co-expressed with GFP::NLS in live animals. Surprisingly, we found that the incorporation efficiency into mNB was lower than for eNB. We therefore used the NMD deficient *smg-6(ok1794)* deletion background to assay mNB variants. Furthermore, we took advantage of higher solubility of the K-NPY dipeptide and grew animals for NPY incorporation on 2mM K-NPY plates. Upon imaging worms grown on 0.1mM ONBY and 2mM K-NPY, we found that both mNB^Y116ONBY^ and mNB^Y116NPY^ localised to the nucleus prior to uncaging (Figure 6B). In neither case was a significant change in localisation observed following uncaging (Figure 6C). This result indicates that, as we previously observed with eNB, introducing a photocaged tyrosine to the interface was not sufficient to break the nanobody/antigen interaction *in vivo*.

We proceeded to run MD simulations followed by BAlaS scans and found several alanine mutations with effects on GFP binding ranging from +0.02kJ/mol to +17.90kJ/mol (Supplementary Figure 8). We selected two mutations, D98A and D119A (Figure 6D), and introduced them separately into mNB alongside the Y116 amber mutation. We generated transgenic *C. elegans* strains expressing these mNB mutant variants together with the incorporation machinery, and GFP::NLS. We then grew transgenic animals on plates supplemented with 2mM K-NPY to express mNB^Y116NPY, D98A^ or mNB^Y116NPY, D119A^ respectively. We found that mNB^Y116NPY, D98A^ was nuclear before uncaging, with an N/C of 1.71 (± 0.22), indicating that the addition of a D98A alanine mutation was not sufficient to break the interaction (Figure 6B, 6C). In contrast, we found that the mNB^Y116NPY, D119A^ nanobody variant was distributed uniformly throughout the cell and translocated to the nucleus following uncaging, with N/C going from 1.06 (± 0.06) to 1.31 (± 0.11). The D119A mutation in addition to Y116NPY was therefore sufficient to break mNB/GFP binding, while uncaged mNB^D119A^ retained sufficient affinity to bind to GFP.

## Discussion

Nanobodies are small, highly specific protein binders, which have been exploited to develop a vast range of *in vivo* tools for investigating biological processes. Several avenues have been explored to date to develop methods for making nanobody antigen binding inducible. The ability to control binding with light would provide the greatest spatiotemporal control of nanobody/antigen interaction, but progress to date has been unsatisfactory, with problems such as slow kinetics, background activity, and a dependence on blue light. The use of photocaged amino acids to control nanobody/antigen binding may help to overcome these limitations as uncaging is rapid and the photocaging groups are stable in blue light, making their application compatible with commonly used imaging and optogenetic approaches.

In principle, the introduction of photocaged residues into binding interfaces can be used to disrupt the formation of PPI, while removal of the photocaging group will restore the native protein and therefore its native binding ability. A difficulty in using photocaged amino acids for this purpose is that replacement of a native residue with its caged counterpart alone may not be sufficient to entirely break the PPI, especially in intracellular environments, thus resulting in considerable residual binding. Here we provide a solution to this problem by combining rational interface engineering, guided by computational alanine scanning and molecular dynamics simulations, with the site-specific incorporation of photocaged ncAA to design improved variants of two anti-GFP nanobodies for use in cellular environments. The introduction of the photo-caged tyrosines ONBY or NPY into the improved variants abolishes binding to GFP in the intracellular environment within a living animal, and removal of the caging group upon illumination restores binding. Our approach will be applicable to other nanobodies and will also allow the control of the binding interaction through tuning and photo-caging of the target protein.

We establish the use of the photocaged tyrosines ONBY and NPY in *C. elegans* using an enhanced genetic code expansion machinery.^[32,35–38]^ The photocaged tyrosine ONBY has previously been used to photocage nanobody/antigen binding *in vitro* and when exogenously delivered to cells.^[25,26]^ We have found that the introduction of an ONBY residue into the GFP binding interface of eNB is not sufficient to break binding when photocaged nanobody and antigen are expressed together in an intracellular environment. It is not entirely surprising that developing a photo-controllable nanobody for use inside a living cell requires additional engineering steps beyond the simple introduction of a photocaged amino acid residue. *In vitro* conditions where proteins are in a controlled, dilute buffer often do not fully replicate conditions encountered in the crowded interior of cells where the environment is composed of a large variety of tightly packed macromolecules.^[45]^ Factors such as nanobody concentration, the intracellular “incubation” time (which begins from the time of expression and may therefore be longer than that of *in vitro* experiments), and the effects of macromolecular crowding may all contribute to the failure of the photocaged tyrosines to break binding *in vivo*.

We have encountered this phenomenon for both of the anti-GFP nanobodies we tested, suggesting that the introduction a photocaged amino acid alone may not be a feasible approach for the development of photocaged nanobodies for *in vivo* applications in general. Our data suggest that even nanobodies with reduced binding affinities are competent binders in intracellular environments. Therefore, the exceptionally strong antigen affinity of nanobodies is not a required or even desirable trait when engineering photo-activatable nanobodies for intracellular *in vivo* use as it may result in substantial photocaged nanobody/antigen interaction pre-activation. Indeed, molecular-dynamics simulations of the wild-type and photocaged nanobodies presented here, suggest that the photocaging groups themselves can add stabilising interactions with the antigen that contribute to background binding. This highlights the complex nature of these interactions and demonstrates the value of a rigorous computational approach to engineering interfaces such as these. Furthermore, our data show the importance of *in vivo* validation for *in silico* PPI predictions and *in vitro* PPI measurements.

Our method represents a novel application of Computational Alanine Scanning (CAS) and a rare example of the results being experimentally validated in a system as complex as a living animal. Our *in vivo* localisation-based binding assay provides a simple, effective method to validate computationally predicted interaction hot-spots in physiologically relevant environments. We believe our work demonstrates that *in vivo* validation methods should be considered when designing hot-spot targeted drugs and antibodies

Of the nanobody mutants tested, three out of six resulted in functional photoinducible nanobodies. Interestingly, one of the mutations that failed to break binding, W47A in eNB, was predicted by BAlaS to have the largest contribution to the free-energy of binding. This might indicate a bias in the scoring function used in BAlaS, which results in over weighting of polar residues vs nonpolar residues. BAlaS was found to produce more accurate results when the contributions of polar residues (DERKH) are down-weighted.^[30]^ Our observations indicate that this correction could be too severe. As noted in the Results, BAlaS suggested Y37 as an important residue, which also indicates that CAS could be used to identify candidate residues to photocage. In future, we plan to further develop this pipeline to fully automate the process of selecting the identity and location of the photocaged residue, as well as possible secondary mutations to tune the interaction strength, to make the process of generating light-inducible interactions more accessible to the broader scientific community.

Importantly, by tuning the binding energy through mutations informed by our *in silico* alanine scanning approach, we were able to photocage nanobody/GFP binding using the photocaged tyrosine NPY. NPY is more amenable to *in vivo* applications than ONBY due to its superior uncaging properties, with a red shifted absorption maximum of 365 nm compared to 254 nm for ONBY. Using our approach we were able to generate photo-controllable variants of the anti-GFP enhancer nanobody (eNB), which is used as the basis of a wide range of *in vivo* tools.^[2,11,12,16–18]^ These variants may thus immediately add the ability to use light for the precise spatiotemporal control of these tools, with the photocaged tyrosine that is most amenable to *in vivo* systems, NPY.

## Conclusion

In conclusion, we have developed a generally applicable method for the rational engineering of photo-activatable nanobodies for use *in vivo*. We have shown the feasibility of tuning binding energies through the targeted introduction of alanine mutations into the binding interface as directed by CAS. A general method for the engineering of photocaged nanobodies will allow the addition of precise spatiotemporal control to the large and growing number of existing nanobody based tools, greatly enhancing their power to drive discovery in the large number of model systems in which they are employed.

## Methods

### *C. elegans* maintenance

*C. elegans* strains were maintained under standard conditions unless otherwise stated.^[46,47]^

### Plasmid generation

All plasmids used for transgenesis of *C. elegans* are described in Supplementary Table 1. Plasmids for expression in *C. elegans* were generated using 3 Fragment Multisite Gateway Cloning (Thermo Fisher Scientific).

Details on cloning are listed in Supplementary Table 2. All primers and synthetic genes (gBlocks) were synthesised by IDT. All genes were optimised for expression in *C. elegans* using the online *C. elegans* codon adaptor.^[48]^ PCRs were carried out using Q5 2x Hot-Start Master Mix (New England Biolabs). PCR and digestion products were recovered from agarose gels following electrophoresis using the Zymogen Gel Recovery Kit (Zymo Research). DNA assembly was performed using NEBuilder 2x Master Mix (New England Biolabs). Standard ligation of restriction enzyme digest products was performed using the Roche Rapid Ligation Kit (Merck Life Sciences). All reagents were used in accordance with the manufacturer’s specifications.

Entry and expression plasmids were grown in NEB-5α cells (New England Biolabs). Destination plasmids were grown in One Shot *ccdB* survival cells (Thermo Fisher Scientific). Transformations were performed in accordance with the manufacturer’s specifications. Transformed cells were grown overnight on TB agar containing the appropriate selection antibiotics. Plasmids were recovered from bacteria using the QIAprep Spin Miniprep kit (Qiagen).

### *C. elegans* transgenesis

Transgenic *C. elegans* strains were generated by biolistic bombardment into either the N2 or *smg-6(ok1794)* genetic backgrounds, with Hygromycin B resistance used as the selection marker as previously described.^[49]^ Spermidine (Merck Life Sciences) was used to precipitate DNA onto gold particles of 0.3-3µm diameter (ChemPur). 900psi rupture discs (Bio-Rad) and macro carrier discs (Inbio Gold) were used in a PDS1000/He Biolistic Particle Delivery System (Biorad). Transgenic strains were maintained on NGM agar supplemented with 0.3mg/mL Hygromycin B (Formedium). All strains used in this study are described in Supplementary Table 3.

### Feeding of photocaged amino acids

The amino acid NPY and the dipeptide K-NPY were custom synthesised (NewChem Technologies), and ONBY was purchased (Fluorochem). K-NPY was dissolved in water and added to molten NGM agar to make the desired concentration. NPY and ONBY were dissolved in 5M NaOH before addition to molten NGM agar, followed by neutralisation by HCl.

Before feeding with ncAAs, worms were grown on NGM agar plates seeded with a lawn of *E. coli* OP50 bacteria. Animals were grown until the food was depleted and the population contained a large number of age synchronised L1 larvae. Synchronised populations of L1 animals were washed off starved plates using M9 buffer and transferred to ncAA-NGM agar plates. 40uL of dissolved freeze-dried OP50 (LabTIE) was added to the plates as food. Animals were grown on ncAA plates at 20°C for 24-48 hours before uncaging followed by imaging or western blotting.

### Uncaging

Animals were uncaged under a 365nm LED with the output set to 30% (10mW/cm^2^). Animals were transferred from ncAA-NGM agar plates to NGM-only plates before uncaging. Animals were imaged immediately after uncaging for the 640s timepoints, for shorter illumination times the animals were imaged 640s after the start of illumination.

### Worm lysis and Western blotting

Synchronised populations were grown on ncAA-NGM agar plates at 20°C for 24-48 hours, then washed off plates using M9 buffer (supplemented with 0.001% Triton-X100 to prevent animals from sticking to pipette tips). Worms were settled by gravity, the supernatant was removed, and worms were resuspended in lysis buffer at a 1:2 volume ratio of worms to lysis buffer. The lysis buffer consisted of 4xLDS loading buffer supplemented with reducing agent (Thermo Fisher Scientific) for a 4:1 ratio of 4xLDS to reducing agent. Lysis was performed by a freeze/thaw cycle of overnight freezing at −80°C followed by 15 min incubation at 95°C while shaking.

Samples were run on precast Bolt 4 to 12% gels (Thermo Fisher Scientific) for 18 min at 200V. Proteins were transferred from the gel onto a nitrocellulose membrane using an iBlot2 device (Thermo Fisher Scientific).

After transfer, the membrane was blocked with 5% milk powder in PBST (PBS supplemented with 0.1% Tween-20) for 1h at room temperature. Incubation with primary antibodies was carried out in PBST + 5% milk powder at 4C overnight. Blots were washed 4 x 5 minutes with PBST + 5% milk powder before incubation with secondary antibody for 1h at room temperature.

The primary antibodies used were mouse anti-GFP (clones 7.1 and 13.1) (Roche) at a dilution 1:5000 for SGR57 and SGR58, rat anti-HA clone 3F10 (Roche) at a dilution of 1:1000 for SGR57, and mouse anti-mCherry-Tag Monoclonal (Elabscience) at a dilution of 1:1000 for SGR58. The secondary antibodies used were Horse anti-mouse IgG HRP (Cell Signalling Technology) at a dilution of 1:5000 for anti-GFP and 1:3000 for anti-mCherry-Tag Monoclonal, and Goat anti-Rat IgG (H+L) HRP (Thermo Fisher Scientific) 1:5000. Pierce ECL Western Blotting Substrate (Thermo Fisher Scientific) or SuperSignal West Femto chemiluminescent Substrate (Thermo Fisher Scientific) were used as detection agent.

### Imaging

All microscopy imaging was carried out on a Zeiss M2 imager. Animals were mounted on 2.5% agar pads on glass slides. To immobilise, animals were picked into drop of 50mM NaN_3_ (Thermo Fisher Scientific) on the pad. The NaN_3_ was diluted from a 100mM stock using M9. Animals picked into the drop were left for 3-4 minutes before imaging.

### Measurement of mCherry nuclear/cytoplasmic ratio

Measurement of mCherry nuclear/cytoplasmic ratio in animals expressing photocaged nanobody variants was performed using ImageJ software. Nuclear average brightness was measured in a region of interest within the nucleus avoiding interference from the nucleolus, from which nanobody::mCherry fusions appeared excluded. Cytoplasmic average was measured from the cytoplasmic area proximal to the nuclei, as per the method described by Kelley and Paschal.^[50]^ The ImageJ “threshold” function was used to generate regions of interest encompassing the nuclei in the GFP channel. These “small” ROIs were used to generate a mask which was then enlarged by applying the ImageJ “dilate” function three times to generate the “large” ROIs which extended beyond the nucleus. The area and average brightness of these “small” and “large” ROIs were measured, and cytoplasmic brightness was acquired by the following formulae:

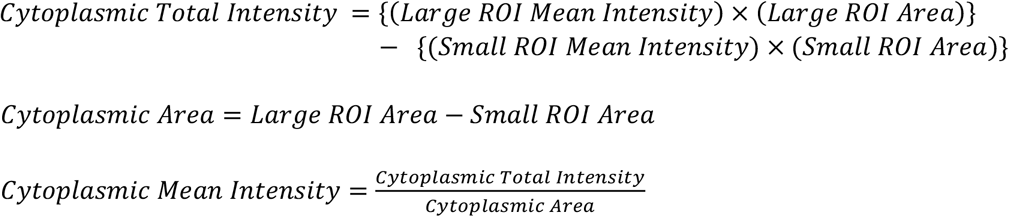

### Statistical analysis

Statistical significance of subcellular localisation results was determined using a two-tailed Welch’s T-test. Regressions were fit using a one-phase linear decay interpolation. Calculations were performed using GraphPad Prism version 9.0.2 for Windows (GraphPad Software).

### Parameterisation of photocaged amino acids for simulation

To prepare input files for MD simulations of eNB^Y37ONBY^ and eNB^Y37NPY^ it was necessary to generate parameters for the ONBY and NPY caging groups. These parameters were derived from the Generalised Amber Force Field (GAFF)^[42]^ and supplemented with quantum mechanical parameters for the nitro-group.^[43]^ A detailed description of the process, including files, code, and Python environment used to generate the input files for these simulations, can be found at https://github.com/wells-wood-research/oshea-j-interface-engineering-2021.

### Molecular dynamics

Files were prepared for molecular dynamics using AmberTools18^[42]^ and Open Babel.^[51]^ Molecular dynamics simulations were performed using OpenMM.^[44]^ The non-bonding interactions were modelled by PME, and the cut-off for non-bonding interactions was 1nm. Bonds involving hydrogen atoms were constrained in length. Simulations were run at 1bar and 300K. Pressure was maintained by a Monte Carlo barostat and temperature was maintained by the Langevin integrator. The frictional constant for the Langevin integrator was 1ps^-1^. A timestep of 2fs was used. A detailed description of the simulations, including the files, code, and Python environment used to perform them, can be found at https://github.com/wells-wood-research/oshea-j-interface-engineering-2021.

### Alanine scanning

Alanine scanning was performed using the open-source program BUDE Alanine Scan. A detailed description of the process, including the environment, files, and code used, can be found at https://github.com/wells-wood-research/oshea-j-interface-engineering-2021.

## Acknowledgements

We thank Maria Doitsidou, Ted Hupp, Kathrin Lang, and Lloyd Davis for helpful suggestions and input on the manuscript, and the members past and present of the Sessions Lab at the University of Bristol for their guidance on using BUDE Alanine Scan.

We thank the European Research Council (ERC-StG-679990), the Wellcome-Trust University of Edinburgh Institutional Strategic Support Fund ISSF2, the Royal Society, and the Muir Maxwell Epilepsy Centre for funding to S.G.; CWW is supported by an Engineering and Physical Sciences Research Council Fellowship (EP/S003002/1) and by the Wellcome-Trust University of Edinburgh Institutional Strategic Support Fund ISSF3.

Some strains were provided by the Caenorhabditis Genetics Centre for strains, funded by NIH Office of Research Infrastructure Programs (P40OD010440)

