## Supplementary Figures and Tables for "Generation of photocaged nanobodies for *in vivo* applications using genetic code expansion and computationally guided protein engineering"

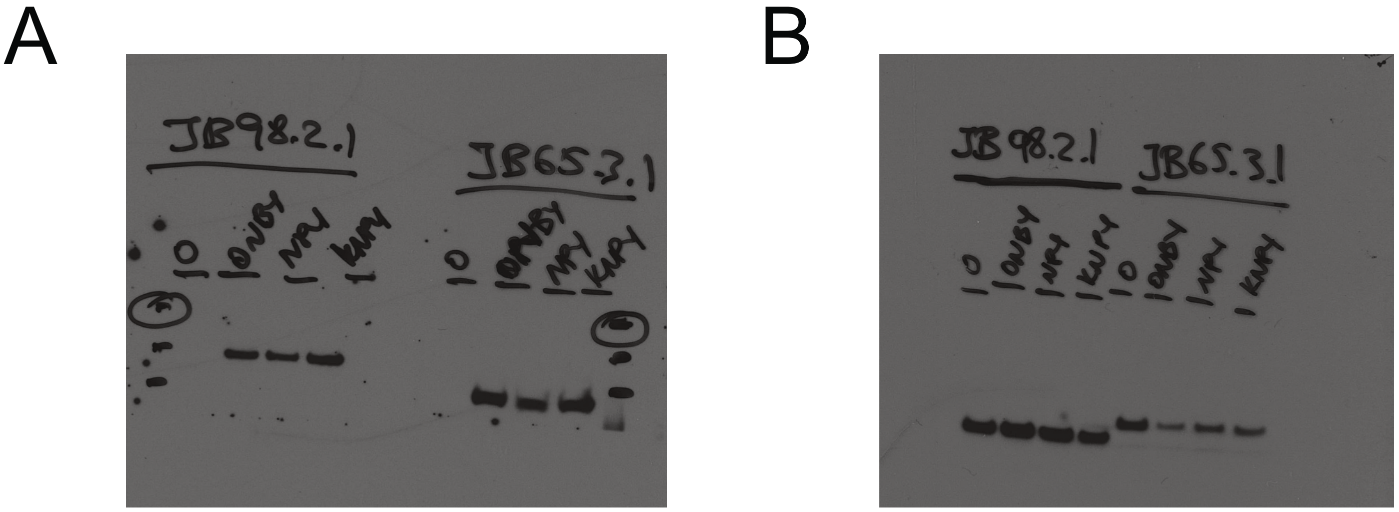


**Supplementary Figure 1** Full scans of Western blots shown in Figure 2C ("JB98.2.1") and Figure 3D ("JB65.3.1"). After transfer, membranes were cut in half, the top half with the ncAA containing proteins, was probed with either anti-HA or anti-mCherry antibodies. The bottom half was probed with anti-GFP and used as a loading control. **A. Left:** Anti-HA Western blots performed on lysates of strain SGR57 ("JB98.2.1"). Animals were grown in the absence of non-canonical amino acid or in the presence of 0.1mM ONBY, NPY, or K-NPY. **Right:** Anti-mCherry Western blots performed on lysates of strain SGR58 ("JB65.3.1"). Animals were grown in the absence of non-canonical amino acid or in the presence of 0.1mM ONBY, NPY, or K-NPY. **B.** anti-GFP Western blots of the membranes shown in (A).

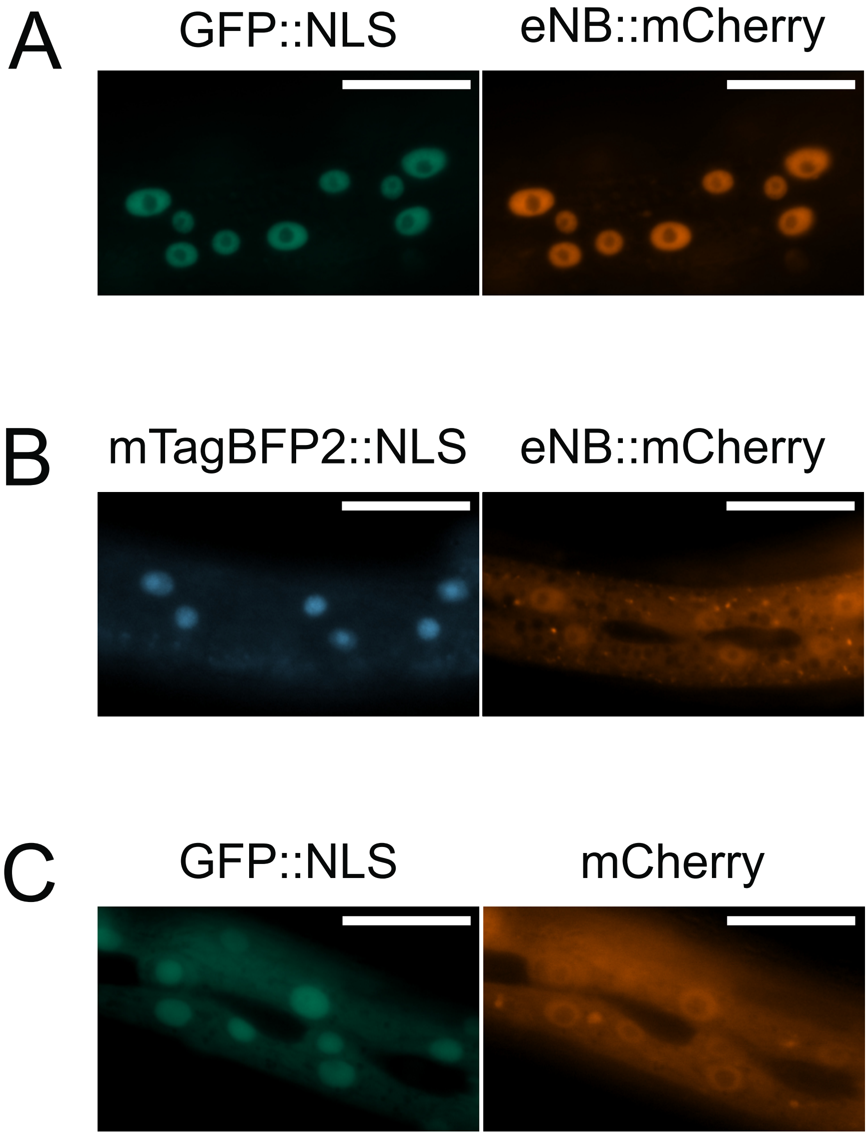


**Supplementary Figure 2 A.** Fluorescence images of *C.elegans* expressing wildtype eNB fused with mCherry and nuclear GFP. **B.** Fluorescence images of *C.elegans* expressing wildtype eNB fused with mCherry and nuclear mTagBFP2. **C.** Fluorescence images of *C.elegans* expressing mCherry and nuclear GFP. All Scale bars 20μm.

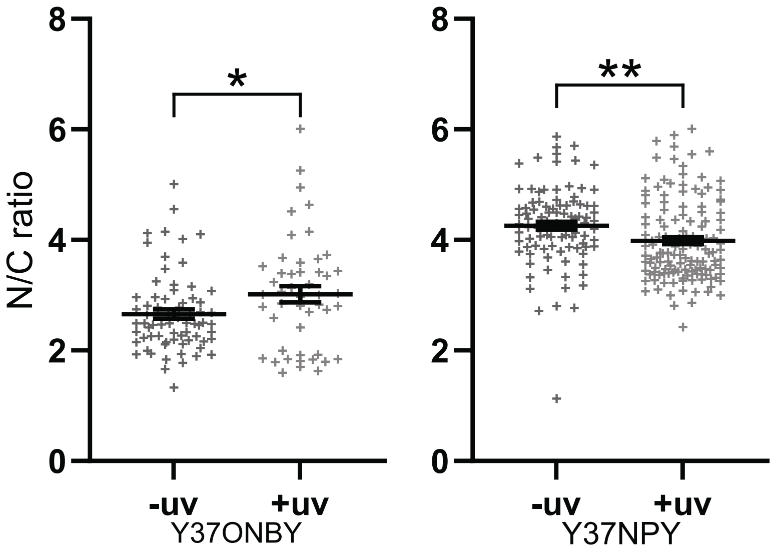


**Supplementary Figure 3** Quantification of eNB::mCherry nuclear/cytoplasmic ratio for eNB with mutations Y37ONBY or Y37NPY before and after uncaging with 365nm. Data are presented as measurements of individual cells and mean ± SEM. Measurements were taken from 7-10 animals per condition. * $p<0.05$; ** $p<0.01$

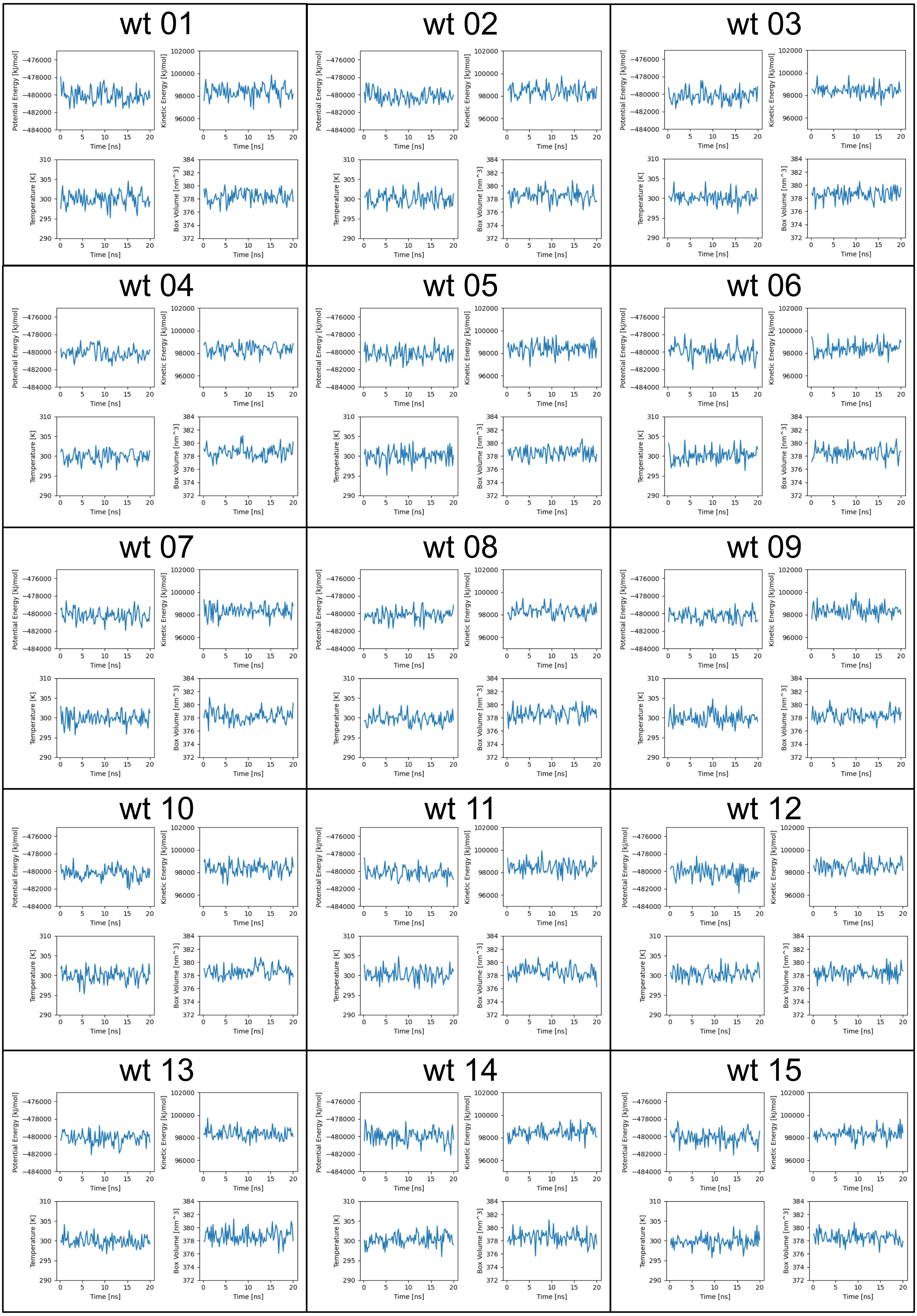


**Supplementary Figure 4** Plots of potential energy, kinetic energy, temperature, and box volume of molecular-dynamics simulations for $eNB^{wt}$, $eNB^{Y37ONBY}$, and $eNB^{Y37NPY}$.

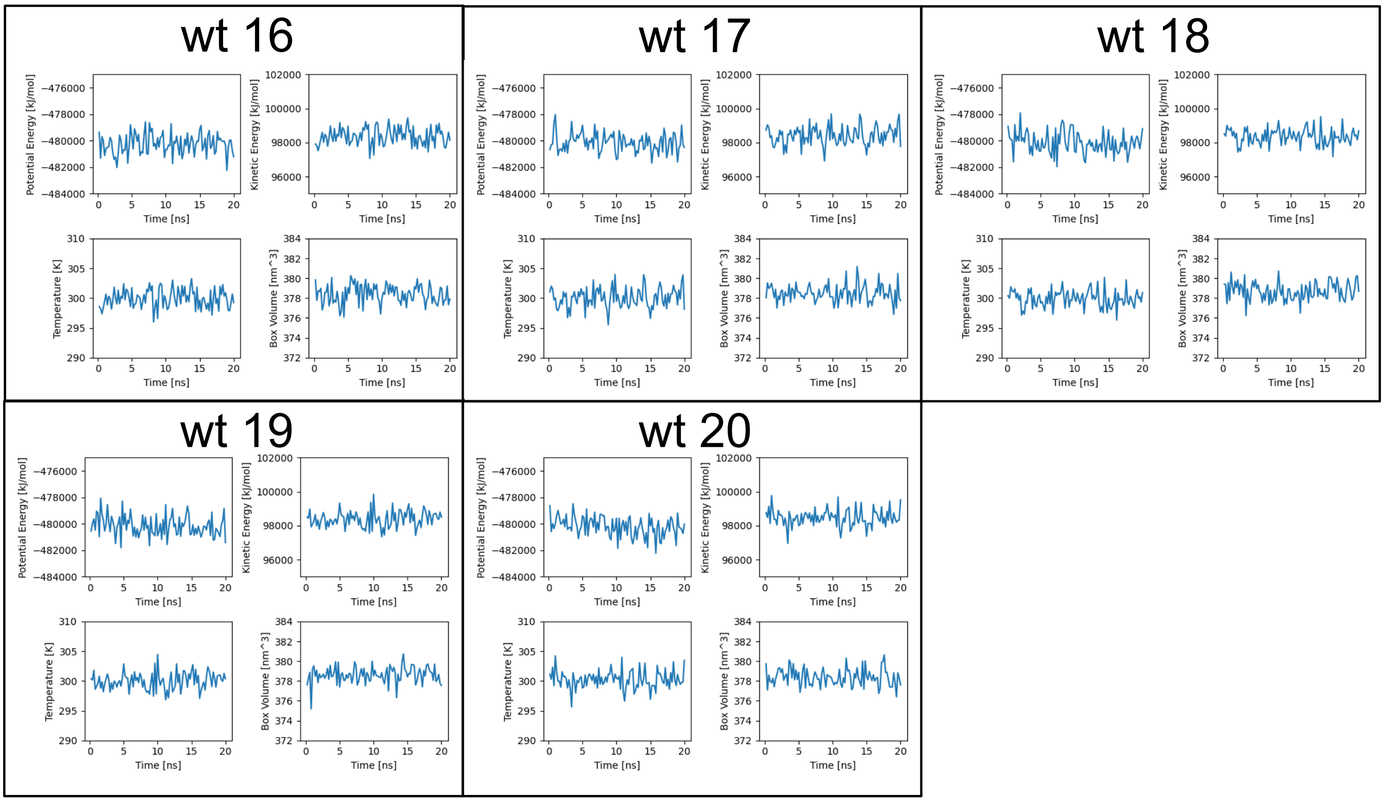


**Supplementary Figure 4 continued.**

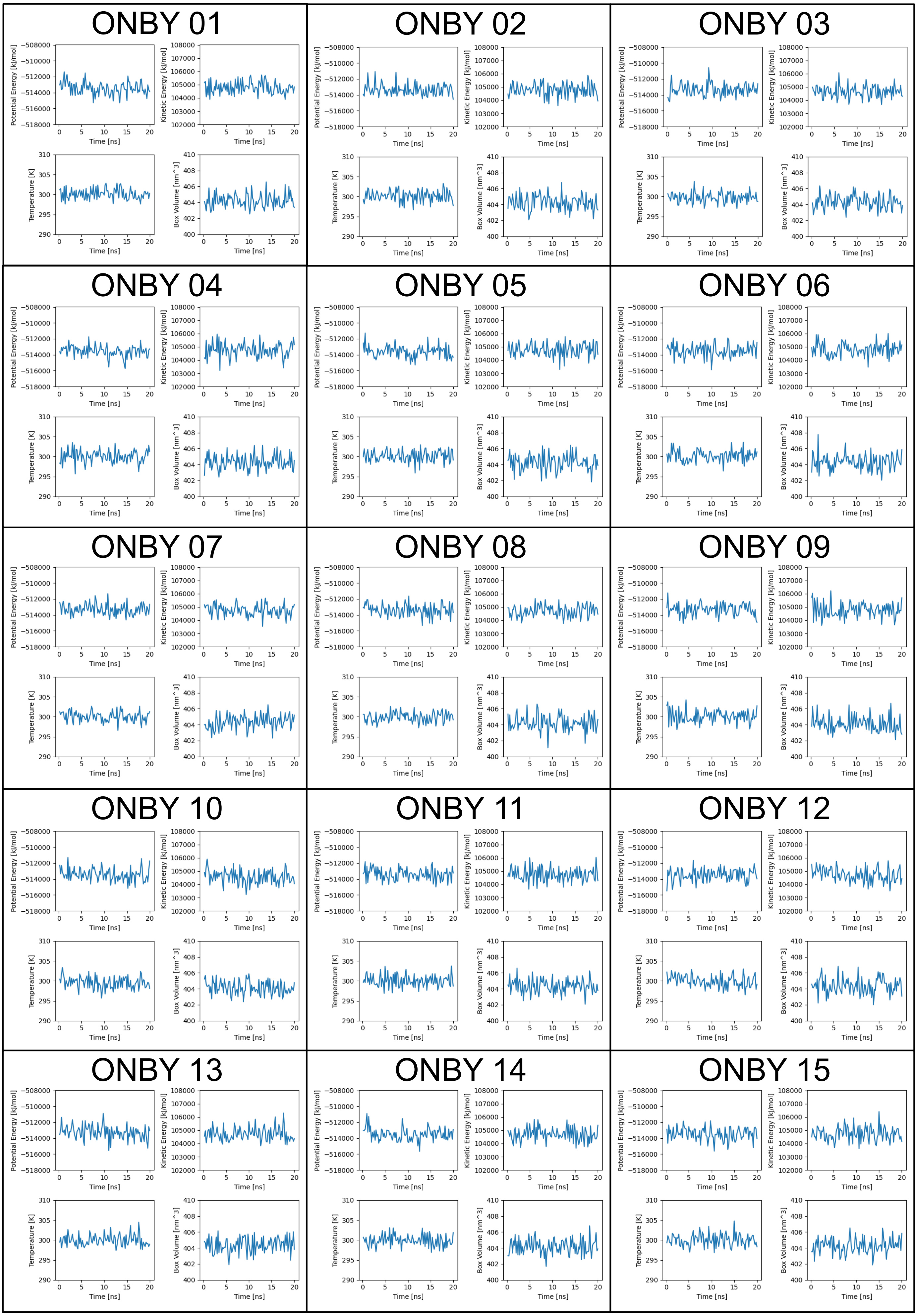


**Supplementary Figure 4 continued.**

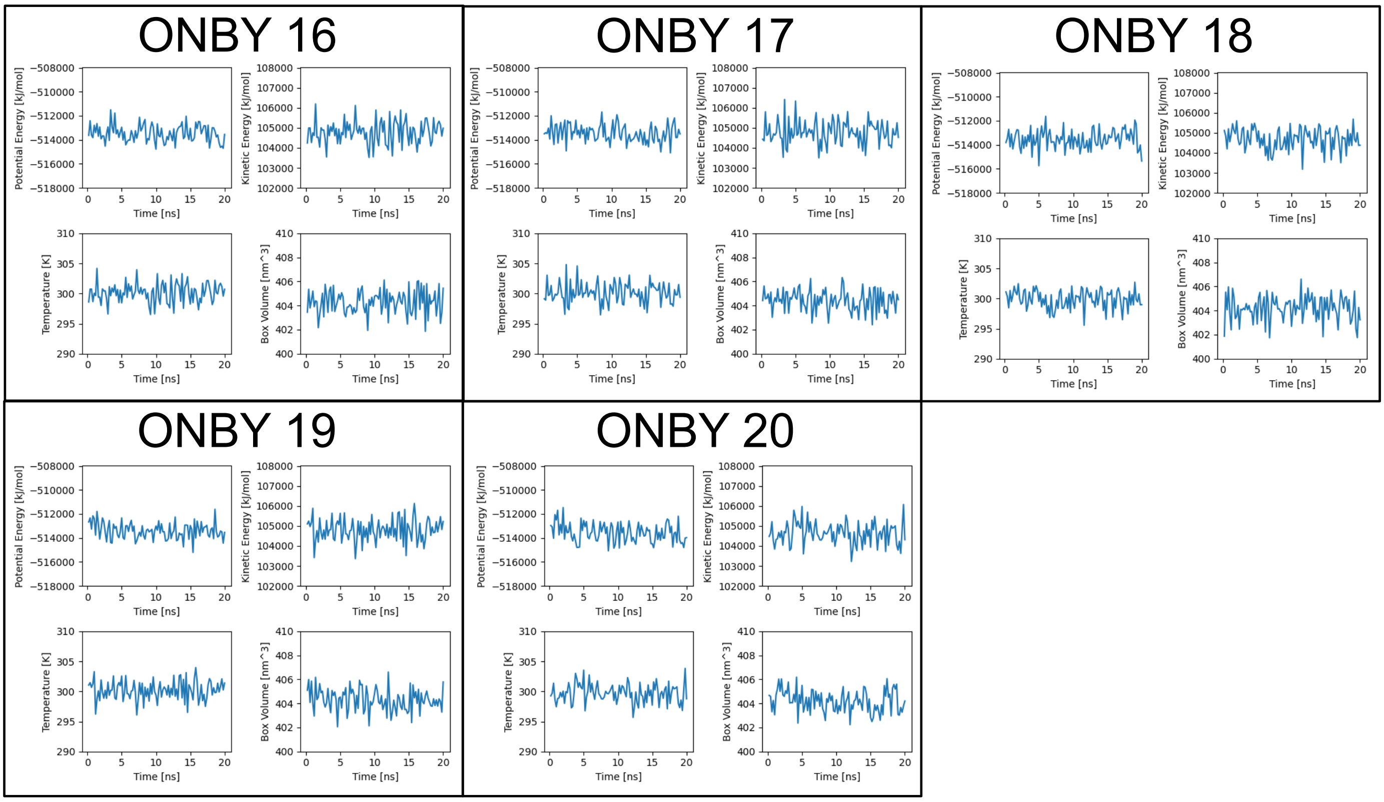


**Supplementary Figure 4 continued.**

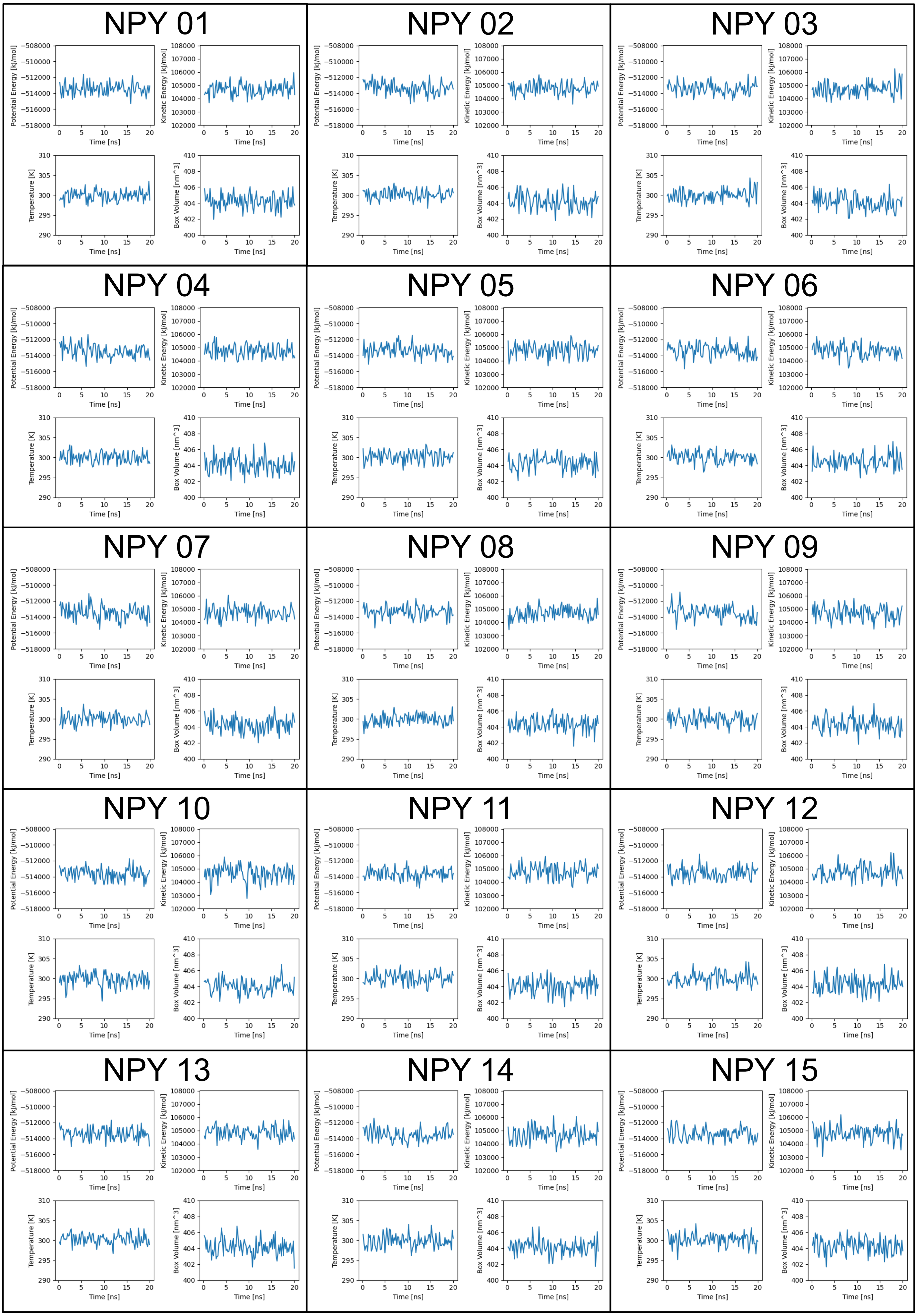


**Supplementary Figure 4 continued.**

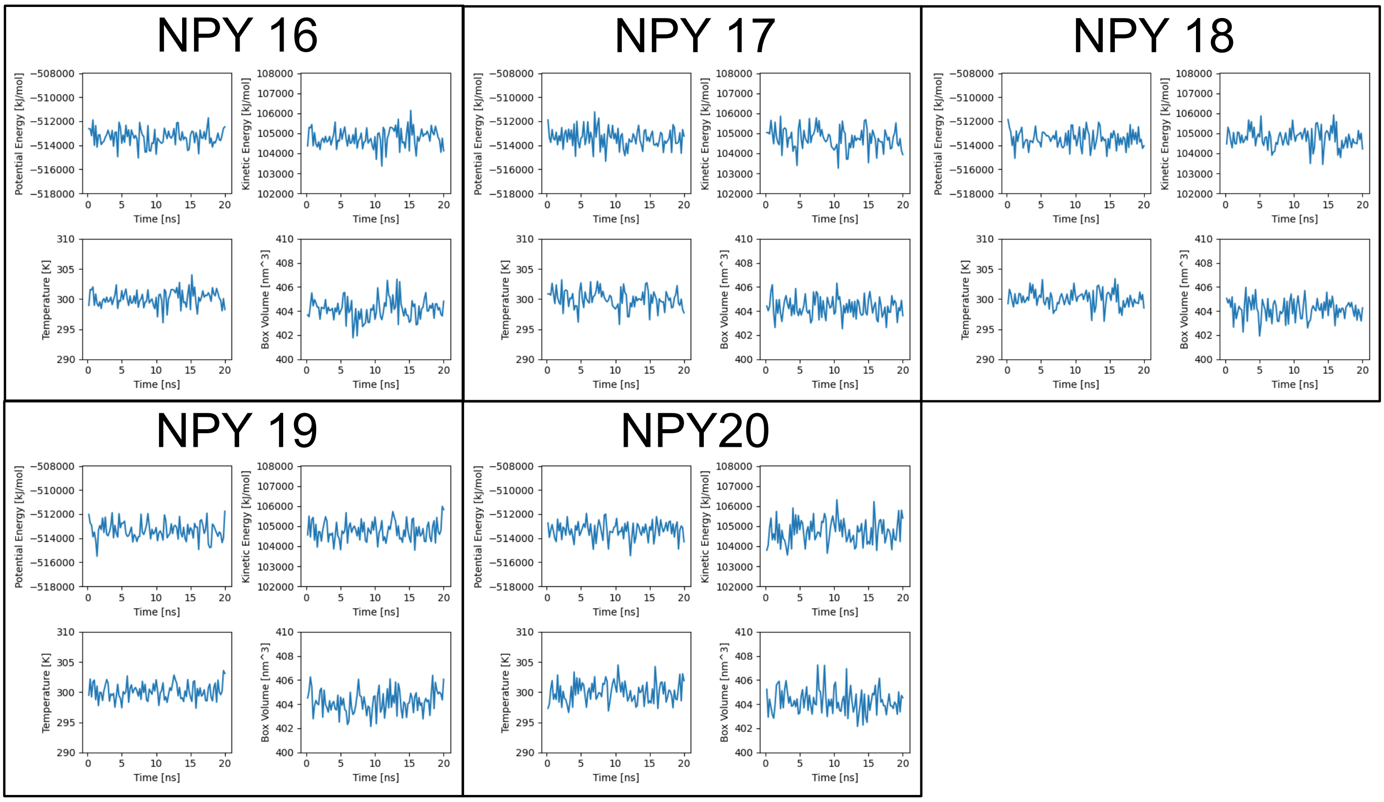


**Supplementary Figure 4 continued.**

A

| Residue number | Residue name | ΔΔG (kJ/mol) |
|---|---|---|
| 31 | ARG | 0.01 |
| 32 | TYR | 0.30 |
| 33 | SER | 1.53 |
| 34 | MET | 0.02 |
| 35 | ARG | 6.69 |
| 37 | TYR | 4.88 |
| 44 | GLU | 2.16 |
| 45 | ARG | 0.51 |
| 46 | GLU | 1.59 |
| 47 | TRP | 9.01 |
| 51 | MET | 0.21 |
| 52 | SER | 0.25 |
| 53 | SER | 0.43 |
| 56 | ASP | 0.41 |
| 57 | ARG | 3.67 |
| 59 | SER | 2.51 |
| 60 | TYR | 0.17 |
| 61 | GLU | 1.18 |
| 62 | ASP | 0.80 |
| 70 | ILE | 0.19 |
| 97 | ASN | 0.51 |
| 98 | VAL | 0.04 |
| 99 | ASN | 3.85 |
| 100 | VAL | 0.04 |
| 102 | PHE | 9.62 |
| 103 | GLU | 6.28 |
| 104 | TYR | 0.42 |
| 105 | TRP | 3.89 |

B

| Residue number | Residue name | ΔΔG (kJ/mol) |
|---|---|---|
| 102 | PHE | 9.62 |
| 47 | TRP | 9.01 |
| 35 | ARG | 6.69 |
| 103 | GLU | 6.28 |
| 37 | TYR | 4.88 |
| 105 | TRP | 3.89 |
| 99 | ASN | 3.85 |
| 57 | ARG | 3.67 |
| 59 | SER | 2.51 |
| 44 | GLU | 2.16 |
| 46 | GLU | 1.59 |
| 33 | SER | 1.53 |
| 61 | GLU | 1.18 |
| 62 | ASP | 0.80 |
| 45 | ARG | 0.51 |
| 97 | ASN | 0.51 |
| 53 | SER | 0.43 |
| 104 | TYR | 0.42 |
| 56 | ASP | 0.41 |
| 32 | TYR | 0.30 |
| 52 | SER | 0.25 |
| 51 | MET | 0.21 |
| 70 | ILE | 0.19 |
| 60 | TYR | 0.17 |
| 98 | VAL | 0.04 |
| 100 | VAL | 0.04 |
| 34 | MET | 0.02 |
| 31 | ARG | 0.01 |

C

```
  1 QVQLVESGGALVQPGGSLRLSCAASGFPVNRYSMRWYRQAPGKEREWVAG
 51 MSSAGDRSSYEDSVKGRFTISRDDARNTVYLQMNSLKPEDTAVYYCNVNV
101 GFEYWGQGTQVTVS
```

**Supplementary Figure 5 A.** Results of BUDE Alanine Scan analysis of eNB/GFP interaction ordered by residue number. The number and identity of eNB residues are given along with the change in ΔG (ΔΔG) that occurs when the residue is mutated to alanine with respect to the wild-type ΔG. Residues selected for experimental analysis are highlighted in blue. The tyrosine selected for photocaging is highlighted in red. **B.** Results of BUDE Alanine Scan analysis of eNB/GFP interaction ordered by ΔΔG. **C.** Amino acid sequence of eNB with relevant residues labeled in blue and Y37 labeled in red.

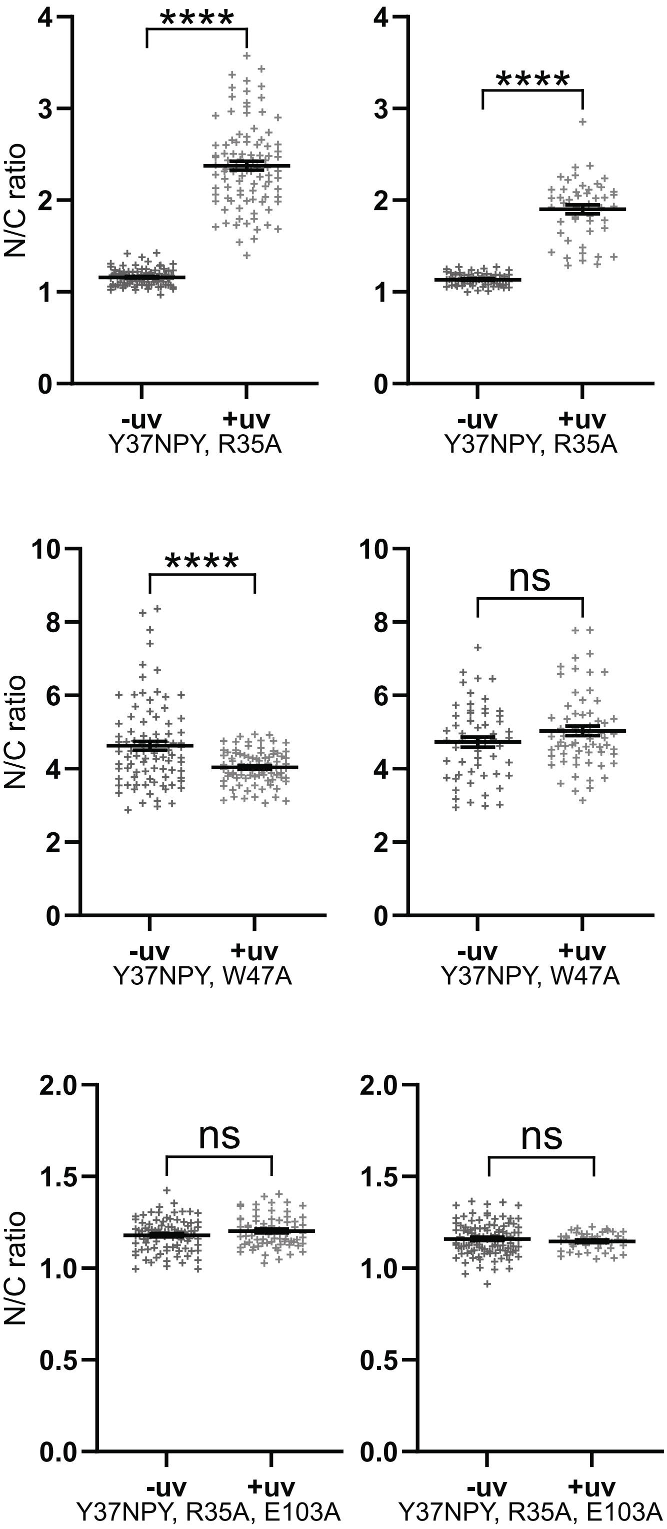


**Supplementary Figure 6** Repeats of quantifications of eNB::mCherry nuclear/cytoplasmic ratio for eNB variants. Data are presented as measurements of individual cells and mean ± SEM. Measurements were taken from 7-10 animals per condition. ns p>0.05; **** p<0.0001

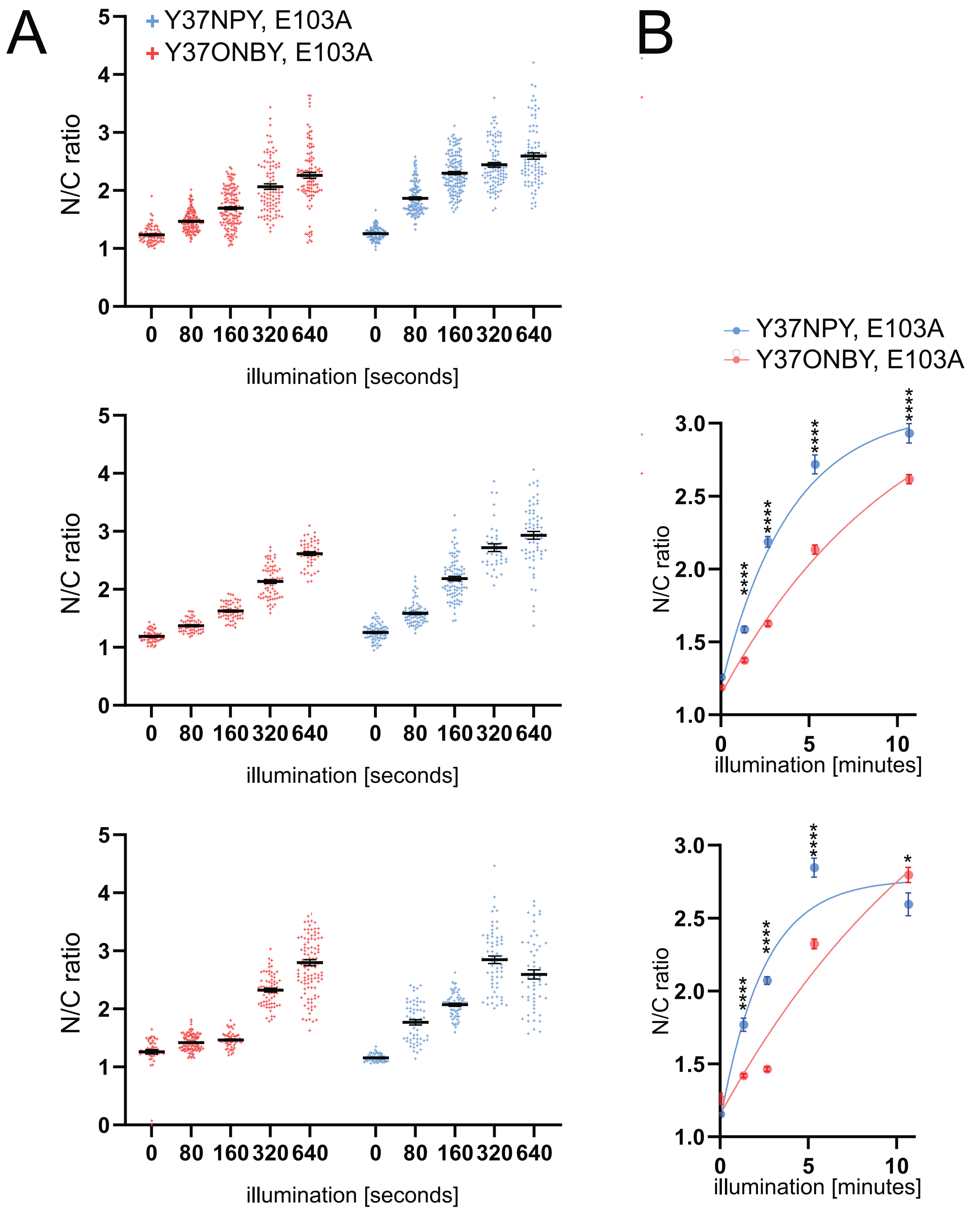


**Supplementary Figure 7. A.** Quantification of eNB::mCherry nuclear/cytoplasmic ratio for eNB$^{Y37ONBY, E103A}$ and eNB$^{Y37NPY, E103A}$ subjected to a range of 365nm illumination times. Data are presented as individual cell measurements and mean ± SEM. Measurements were taken from 7-10 animals per condition. **B.** Quantification of mCherry nuclear/cytoplasmic ratio of eNB$^{Y37ONBY, E103A}$ and eNB$^{Y37NPY, E103A}$ subjected to a range of 365nm illumination times. Data are presented as mean ± SEM. Measurements were taken from 7-10 animals per condition.* $p < 0.1$; ** $p < 0.01$; **** $p < 0.0001$. Top ONBY 000s vs NPY 000s $p = 0.0011$; bottom ONBY 000s vs NPY 000s $p = 0.0087$.

A

| Residue number | Residue name | ΔΔG (kJ/mol) |
|---|---|---|
| 37 | PHE | 0.15 |
| 43 | LYS | 0.02 |
| 44 | GLU | 0.58 |
| 46 | GLU | 0.13 |
| 47 | LEU | 0.32 |
| 50 | ASN | 0.05 |
| 52 | LEU | 0.08 |
| 58 | THR | 0.19 |
| 64 | LYS | 0.60 |
| 98 | ASP | 6.88 |
| 101 | THR | 3.97 |
| 103 | LEU | 7.63 |
| 105 | TYR | 17.90 |
| 106 | VAL | 0.74 |
| 109 | VAL | 1.65 |
| 111 | LEU | 0.42 |
| 115 | ASP | 0.68 |
| 116 | TYR | 9.85 |
| 117 | VAL | 3.71 |
| 118 | MET | 0.11 |
| 119 | ASP | 6.35 |
| 120 | TYR | 0.18 |
| 121 | TRP | 0.03 |

B

| Residue number | Residue name | ΔΔG (kJ/mol) |
|---|---|---|
| 105 | TYR | 17.90 |
| 116 | TYR | 9.85 |
| 103 | LEU | 7.63 |
| 98 | ASP | 6.88 |
| 119 | ASP | 6.35 |
| 101 | THR | 3.97 |
| 117 | VAL | 3.71 |
| 109 | VAL | 1.65 |
| 106 | VAL | 0.74 |
| 115 | ASP | 0.68 |
| 64 | LYS | 0.60 |
| 44 | GLU | 0.58 |
| 111 | LEU | 0.42 |
| 47 | LEU | 0.32 |
| 58 | THR | 0.19 |
| 120 | TYR | 0.18 |
| 37 | PHE | 0.15 |
| 46 | GLU | 0.13 |
| 118 | MET | 0.11 |
| 52 | LEU | 0.08 |
| 50 | ASN | 0.05 |
| 121 | TRP | 0.03 |
| 43 | LYS | 0.02 |

C

```
  1 DVQLQESGGGSVQAGGSLRLSCAASGDTFSSYSMAWFRQAPGKECELVSN
 51 ILRDGTTTYAGSVKGRFTISRDDAKNTVYLQMVNLKSEDTARYYCAADSG
101 TQLGYVGAVGLSCLDYVMDYWGKGTQVTVS
```

**Supplementary Figure 8 A.** Results of BUDE Alanine Scan analysis of mNB/GFP interactions ordered by residue number. The number and identity of mNB residues are given along with the change in ΔG (ΔΔG) that occurs when the residue is mutated to alanine with respect to the wild-type ΔG. Residues selected for experimental analysis are highlighted in blue. The tyrosine selected for photocaging is higlighted in red. **B.** Results of BUDE Alanine Scan analysis of mNB/GFP interaction ordered by ΔΔG. **C.** Amino acid sequence of mNB with the relevant residues highlighted in blue and Y116 in red.

### Supplementary Table 1 – Expression plasmids

| Name | Description | Source plasmids | | | |
|---|---|---|---|---|---|
| | | Destination | P4-P1r | 221 | P2r-P3 |
| JOS97 | Psur-5::HuPKIαNES::MmNPYRS | IR98 | SE72 | KB125 | SG606 |
| JOS77 | Psur-5::mCherry::SL2::GFP::EGL-13NLS | pDEST | SE72 | IR186 | LD343 |
| JOS233 | Psur-5::enhancer::mCherry::SL2::EGL-13NLS::BFP | JOS163 | SE72 | JOS4 | JOS164 |
| JOS137 | Psur-5::enhancer(Y37am, R35A)::mCherry::SL2::GFP::EGL-13NLS | JOS13 | SE72 | JOS117 | JOS19 |
| JOS145 | Psur-5::enhancer(Y37am, E101A)::mCherry::SL2::GFP::EGL-13NLS | JOS13 | SE72 | JOS125 | JOS19 |
| JOS94 | Psur-5::enhancer(Y37am)::mCherry::SL2::GFP::EGL-13NLS | JOS13 | SE72 | JOS41 | JOS6 |
| JOS52 | Psur-5::enhancer::mCherry::SL2::GFP::EGL-13NLS | JOS13 | SE72 | JOS4 | JOS6 |
| JOS140 | Psur-5::enhancer(Y37am, R35A, E103A)::mCherry::SL2::GFP::EGL-13NLS | JOS13 | SE72 | JOS120 | JOS19 |
| JOS142 | Psur-5::enhancer(Y37am, W47A)::mCherry::SL2::GFP::EGL-13NLS | JOS13 | SE72 | JOS122 | JOS19 |
| SG88 | Prps-0::GFP(am)::mCherry::HA::EGL-13NLS | [34] | | | |
| JOS229 | Psur-5::minimiser(Y116am)::mCherry::HA::SL2::GFP::EGL-13NLS | JOS13 | SE72 | JOS40 | JOS164 |
| JOS230 | Psur-5::minimiser(Y116am, D98A)::mCherry::HA::SL2::GFP::EGL-13NLS | JOS13 | SE72 | JOS180 | JOS164 |
| JOS232 | Psur-5::minimiser(Y116am, D119A)::mCherry::HA::SL2::GFP::EGL-13NLS | JOS13 | SE72 | JOS182 | JOS164 |

### Supplementary Table 2.1 – Destination Vectors

| Name | Description | Source |
|---|---|---|
| IR98 | pDEST R4-R3 Prps-0::HygR | [49] |
| JOS13 | pDEST R4-R3 SL2::GFP-NLS::let-858 3' UTR | Made from LD343 and SE114. SL2::GFP-NLS-let858 was amplified from LD343. SE114 was digested with BglII and the amplified construct was assembled with the larger digestion product using NEBuilder.<br>Primers used for amplification:<br>J92 - gibson::SL2 F:<br>CATTTTACGTTTCTCGTTCAACTTTATTATACATAGTTGAgatctAGATCTGCTGTCTCATCCTACTTTCAC<br>J93 - let858::gibson R:<br>TCCCAGTCACGACGTTGTAAAACGACGGCCAGTGAATTAagatcTagatcTATACGGATTCGCATTTGCCA |
| SE114 | pDEST R4-R3 II BglII::let-858 3' UTR::BglII | Made from SG606 and pDESTII R3-R4. pDESTII R3-R4 was opened by PCR amplification. let-858 was amplified from SG606. Both amplified products were digested with BglII, then ligated together.<br>Primers for pDESTII R3-R4 amplification:<br>336: CATAGTTGAgatctTAATTCACTGGCCGTCGTTTTACAAC<br>337: TGAATTAagatcTCAACTATGTATAATAAAGTTGAACGAGAAACG<br>Primers for SG606 amplification:<br>339:<br>CATTTTACGTTTCTCGTTCAACTTTATTATACATAGTTGAgatctCGTGAAGTGGAATCGGATGATC<br>340:<br>TCCCAGTCACGACGTTGTAAAACGACGGCCAGTGAATTAagatcTATACGGATTCGCATTTGCCAAG |
| SG72 | pDEST R4-R3 SL2::GFP::let-858 3' UTR | [34] |
| JOS163 | pDEST R4-R3 SL2::EGL13NLS::mTagBFP2::let-858 3' UTR | Made from JOS13 and gBlock gJ3. JOS13 was opened by PCR. The amplified product was recovered and assembled with gJ3 by NEBuilder.<br>Primers for JOS13 opening:<br>582 - let858 F:<br>TAACGTGAAGTGGAATCGGATGATC<br>579 - SL2 R:<br>TTTTTCTACCGGTACAGCAGTTTCC |

### Supplementary Table 2.2 – pENTR P4-P1r Vectors

| Name | Description | Source |
|---|---|---|

| | | |
|---|---|---|
| SE72 | Psur-5 | Amplified from genonimic DNA.<br>189 P sur-5s attB4F:<br>GGGGACAACTTTGTATAGAAAAGTTGCGCAGGCGGTAAACATACGTTG<br>193 P sur-5 attB1R:<br>GGGGACTGCTTTTTTGTACAAACTTGTCTGAAAACAAAATGTAAAGTTCAAAGG |

### Supplementary Table 2.3 – pENTR 221 Vectors

| Name | Description | Source |
|---|---|---|
| ZX297 | Prpr-1::tRNA(C15)::sup-7 short 4x | Made from ZX296. ZX296 was digested with SalI and BamHI and the fragment containing two "Prpr-1 Bt mttRNA Ser C15 sup-7" repeats was recovered. ZX296 was opened by digest with SalI and BglII. These two digest products were ligated together. |
| ZX296 | Prpr-1::tRNA(C15)::up-7 short 2x | Made from SE150. SE150 was digested with SalI and BamHI and the Prpr-1 Bt mttRNA Ser C15 sup-7 fragment was recovered. SE150 was opened by digest with SalI and BglII. These two digest products were ligated together. |
| SE150 | Prpr-1::mttRNA(C15)::sup-7 short | PylT in SG322 was replaced with Bt mttRNA C15 Sequence:<br>GGAAACCTGgTCAgGgAGAcCGAAcGGACTCTAAATCCGTTCAGCCGGGTTcGATTCCCGGGGTTTCCG |
| JOS4 | enhancer::HL | G-block gJ1 was recombined into pDONR 221 by BP recombinase. |
| JOS5 | minimiser::HL | G-block gJ2 was recombined into pDONR 221 by BP recombinase. |
| JOS41 | enhancer(Y37am)::HL | Made from JOS4. JOS4 was amplified with primers that applied an amber stop codon mutation to Y37. The amplified product was recovered and circularised by NEBuilder.<br>J14 - enhY37 F: CGTTACTCCATGCGTTGGTAGCGTCAAGCCCCAGGAAAGG<br>J15 - enhY37 R: CCTTTCCTGGGGCTTGACGCTACCAACGCATGGAGTAACG |
| JOS117 | enhancer(Y37am, R35A)::HL | Made from JOS41. JOS41 was amplified with primers that applied the mutation R35A. The amplified product was recovered and circularised by NEBuilder.<br>J167 - Cele enhancer(Y37am) R35A F: TTACTCCATGGCTTGGTAGCGTC<br>J168 - Cele enhancer(Y37am) R35A R: GACGCTACCAAGCCATGGAGTAA |
| JOS125 | enhancer(Y37am, E103A)::HL | Made from JOS41. JOS41 was amplified with primers that applied the mutation E103A. The amplified product was recovered and circularised by NEBuilder.<br>J175 - Cele enhancer E103A F: CGTCGGATTCGCTTACTGGGGAC<br>J176 - Cele enhancer E103A R: GTCCCCAGTAAGCGAATCCGACG |
| JOS120 | enhancer(Y37am, R35A, E103A)::HL | Made from JOS117. JOS117 was amplified with primers that applied the mutation E103A. The amplified product was recovered and circularised by NEBuilder.<br>J175 - Cele enhancer E103A F: CGTCGGATTCGCTTACTGGGGAC<br>J176 - Cele enhancer E103A R: GTCCCCAGTAAGCGAATCCGACG |
| JOS122 | enhancer(Y37am, W47A)::HL | Made from JOS41. JOS41 was amplified with primers that applied the mutation W47A. The amplified product was recovered and circularised by NEBuilder.<br>J171 - Cele enhancer W47A F: GGAGCGTGAGGCTGTCGCCGGAA<br>J172 - Cele enhancer W47A R: TTCCGGCGACAGCCTCACGCTCC |
| JOS40 | minimiser(Y116am)::HL | Made from JOS5. JOS5 was amplified with primers that applied an amber stop codon mutation to Y116. The amplified product was recovered and circularised by NEBuilder<br>J10 - minY100B F: CTCTCCTGCCTCGACTAGGTCATGGACTACTGGGG<br>J11 - minY100B R: CCCCAGTAGTCCATGACCTAGTCGAGGCAGGAGAG |
| JOS180 | minimiser(Y116am, D98A)::HL | Made from JOS40. JOS40 was amplified with primers that applied the mutation D98A. The amplified product was recovered and circularised by NEBuilder.<br>J255 - Cele minimiser D95 F:<br>CGCCGCCGCTTCCGGAACCCAACTCGGATACGTCGGAGCCG<br>J244 - Cele minimiser(Y100Bam) D95 R:<br>TTCCGGAAGCGGCGGCGCAGTAGTAACGGGCGG |

| | | |
|---|---|---|
| JOS181 | minimise(Y116am, D119A)::HL | Made from JOS40. JOS40 was amplified with primers that applied the mutation D119A. The amplified product was recovered and circularised by NEBuilder.<br>J256 - Cele minimser(Y100Mam) D101A F: CTAGGTCATGGCTTACTGGGGAAAGGGAACCCAAGTCACCG<br>J257 - Cele minimser(Y100Mam) D101A R: TTCCCCAGTAAGCCATGACCTAGTCGAGGCAGGAGAGTCCGACG |
| KB125 | HuPKIαNES::MmNPYRS | Made from LD184. LD184 was opened at the 5' end of the synthetase by PCR, then a primer containing the Hu PKIa NES incorporated at the opening by NEBuilder.<br>Plasmids for amplification:<br>431 MmPylRS NEB R: CATTTTTGCAGCCTGCTTTTTTGTACA<br>430 MmPylRS NEB F: ATGGACAAGAAGCCACTCAACAC<br>Plasmid for NEBuilder:<br>AAAAGCAGGCTGCAAAAATGCTCGCCCTCAAGCTCGCCGGACTCGACATCATGGACAAGAAGCCACTCAA |
| LD184 | MmNPYRS | G-block gL was recombined into pDONR221 by BP recombinase. |
| SG322 | Prpr-1::PylT::sup-7 | P*rpr-1* was amplified from genomic DNA and fused to PylT by overlap extension PCR, 100bp of the sup-7 3' region was also fused to the 3' end of PylT<br>by overlap extension PCR |

### Supplementary Table 2.4 – pENTR P2r-P3 Vectors

| Name | Description | Source |
|---|---|---|
| LD343 | SL2::GFP-NLS::let-858 3'UTR | Made from IR182. NLS-egl13 was attached to the 3' end of GFP, and flanking BP recombinase binding sites were added for recombination into pDONR P2r-P3 vector<br>Plasmids for amplifaction of fragment 1:<br>140 attB2R 5' SL2 GFP F:<br>GGGGACAGCTTTCTTGTACAAAGTGGgctgtctcatcctactttcacctagttaa<br>214 GFP NLS R:<br>TTCGCGTTTTCACTCAGTTTTGTCGGATTCGCTTTTCGTCTACGGCTCATtcctcccttgtagagctcgtccattccg<br>Plasmids for amplifcation of fragment 2:<br>213 GFP NLS F:<br>TCCGACAAAACTGAGTGAAAACGCGAAGAAGCTTGCCAAGGAAGTTGAAAATtaacgtgaagtggaatcggatgatcgacgccg<br>145 attB3 3' GFP let858 R:<br>GGGGACAACTTTGTATAATAAAGTTGatacggattcgcatttgccaag |
| SG606 | let-858 3'UTR | PCR amplified from genomic DNA<br>Plasmids for amplification:<br>725 let-858 3' attB2R:<br>GGGGACCACTTTGTACAAGAAAGCTGGGTATACGGATTCGCATTTGCCAAGC<br>726 let-858 3' attB3R:<br>GGGGACAACTTTGTATAATAAAGTTGATACGGATTCGCATTTGCCAAGC |
| JOS6 | mCherry | Made from SG79. mCherry was amplified from SG79 and flanking BP recombinase binding sites were added for recombination into pDONR P2r-P3.<br>J30 - attR2::mCherry:<br>GGGGACAGCTTTCTTGTACAAAGTGGTAGTCTCAAAGGGTGAAGAAGATAACATGG<br>J31 - mCherry::attL3 R:<br>GGGGACAACTTTGTATAATAAAGTTGTTTACTTATACAATTCATCCATGCCACCTGT |
| JOS19 | mCherry::HisTagx6 | Made from JOS6. mCherry was amplified from JOS6 and an N-terminal 6x His repeat was added. Flanking BP recombinase binding sites were added for recombination into pDONR P2r-P3.<br>J30 - attR2::mCherry:<br>GGGGACAGCTTTCTTGTACAAAGTGGTAGTCTCAAAGGGTGAAGAAGATAACATGG<br>J141 - mCherry::6xHis::attL2 R:<br>GGGGACAACTTTGTATAATAAAGTTGTTTAATGATGATGATGATGATGCTTATACAATTCATCCATGCCACCTGT |

| | | |
|---|---|---|
| JOS164 | mCherry::HisTagx6::HA | Made from JOS19. mCherry::HisTagx6 was amplified from JOS19 and an N-terminal HA tag was added. Flanking BP recombinase binding sites were added for recombination into pDONR P2r-P3.<br>J30 - attR2::mCherry:<br>GGGGACAGCTTTCTTGTACAAAGTGGTAGTCTCAAAGGGTGAAGAAGATAACAT<br>GG<br>J234 - mCherry::His-tag::HA R:<br>CGTAGTCTGGGACGTCGTATGGGTAATGATGATGATGATGATGCTTATACAATTC<br>ATCC |
| SG79 | mCherry::HA | [34] |
| IR182 | SL2::GFP ::let-858 3' UTR | [35] |

### Supplementary Table 2.5 – G-blocks

| Name | Description | Sequence |
|---|---|---|
| gJ1 | attB1::enhancer::HL::attB2<br><br>*C. elegans* optimised enhancer nanobody and glycine-rich linker | ACAAGTTTGTACAAAAAAGCAGGCTaaaaATGGCCCAAGTCCAACTCGTCGAGTCCGGAGGAGCCCTC<br>GTCCAACCAGGAGGATCCCTCCGTCTCTCCTGCGCCGCCTCCGGATTCCCAGTCAACCGTTACTCCATG<br>CGTTGGTACCGTCAAGCCCCAGGAAAGGAGCGTGAGTGGGTCGCCGGAATGTCCTCCGCCGGAGAC<br>CGTTCCTCCTACGAGGACTCCGTCAAGGGACGTTTCACCATCTCCCGTGACGACGCCCGTAACACCGT<br>CTACCTCCAAATGAACTCCCTCAAGCCAGAGGACACCGCCGTCTACTACTGCAACGTCAACGTCGGAT<br>TCGAGTACTGGGGACAAGGAACCCAAGTCACCGTCTCCTCCATCCTCGGAGCCCCATCCGGAGGAGG<br>AGCCACCGCCGGAGCCGGAGGAGCCGGAGGACCAGCCGGACTCATCCACCCAGCTTTCTTGTACAAA<br>GTGGT |
| gJ2 | attB1::minimiser::HL::attB2<br><br>*C. elegans* optimised minimiser nanobody and glycine-rich linker | ACAAGTTTGTACAAAAAAGCAGGCTaaaaATGGCCGTCCAACTCCAAGAGTCCGGAGGAGGATCCGTC<br>CAAGCCGGAGGATCCCTCCGTCTCTCCTGCGCCGCCTCCGGAGACACCTTCTCCTCCTACTCCATGGCC<br>TGGTTCCGTCAAGCCCCAGGAAAGGAGTGCGAGCTCGTCTCCAACATCCTCCGTGACGGAACCACCAC<br>CTACGCCGGATCCGTCAAGGGACGTTTCACCATCTCCCGTGACGACGCCAAGAACACCGTCTACCTCC<br>AAATGGTCAACCTCAAGTCCGAGGACACCGCCCGTTACTACTGCGCCGCCGACTCCGGAACCCAACTC<br>GGATACGTCGGAGCCGTCGGACTCTCCTGCCTCGACTACGTCATGGACTACTGGGGAAAGGGAACCC<br>AAGTCACCGTCTCCATCCTCGGAGCCCCATCCGGAGGAGGAGCCACCGCCGGAGCCGGAGGAGCCG<br>GAGGACCAGCCGGACTCATCTACCCAGCTTTCTTGTACAAAGTGGT |
| gJ3 | EGL-13 NLS mTagBFP2<br><br>*C. elegans* optimised mTagBFP with 5' SL2 overlap and EGL-13 NLS | GGAAACTGCTGTACCGGTAGAAAAAATGAGCCGAAGACGAAAAGCCAACCCAACTAAACTGTCTGAG<br>AACGCCAAGAAGCTCGCCAAGGAGGTCGAGAACATGGTCTCCAAGGGAGAGGAGCTCATCAAGGAG<br>AACATGCACATGAAGCTCTACATGGAGGGAACCGTCGACAACCACCACTTCAAGTGCACCTCCGAGG<br>GAGAGGGAAAGCCATACGAGGGAACCCAAACCATGCGTATCAAGGTCGTCGAGGGAGGACCACTCC<br>CATTCGCCTTCGACATCCTCGCCACCTCCTTCCTCTACGGATCCAAGgtaagtttaaacatatatatactaactaac<br>cctgattatttaaattttcagACCTTCATCAACCACACCCAAGGAATCCCAGACTTCTTCAAGCAATCCTTCCCA<br>GAGGGATTCACCTGGGAGCGTGTCACCACCTACGAGGACGGAGGAGTCCTCACCGCCACCCAAGACA<br>CCTCCCTCCAAGACGGATGCCTCATCTACAACGTCAAGgtaagtttaaacagttcggtactaactaaccatacatattt<br>aaattttcagATCCGTGGAGTCAACTTCACCTCCAACGGACCAGTCATGCAAAAGAAGACCCTCGGATGG<br>GAGGCCTTCACCGAGACCCTCTACCCAGCCGACGGAGGACTCGAGGGACGTAACGACATGGCCCTCA<br>AGCTCGTCGGAGGATCCCACCTCATCGCCAACGCCAAGgtaagtttaaacatgattttactaactaactaatctgatt<br>taaattttcagACCACCTACCGTTCCAAGAAGCCAGCCAAGAACCTCAAGATGCCAGGAGTCTACTACGTC<br>GACTACCGTCTCGAGCGTATCAAGGAGGCCAACAACGAGACCTACGTCGAGCAACACGAGGTCGCCG<br>TCGCCCGTTACTGCGACCTCCCATCCAAGCTCGGACACAAGCTCAACTAACGTGAAGTGGAATCGGAT<br>GATC |

| | | |
|---|---|---|
| gL | attB1::MmNPYRS::attB2<br><br>*Methanosarcina mazei* pyrrolysine aminoacyl tRNA synthetase mutated to recognise photocaged tyrosines and codon optimised for expression in *C. elegans* | GGGGACAAGTTTGTACAAAAAAGCAGGCTATGGACTACAAGGACGACGACGACAAGATGGACAAGA<br>AGCCACTCAACACCCTCATCTCCGCCACCGGACTCTGGATGTCCCGTACCGGAACCATCCACAAGATCA<br>AGCACCACGAGGTCTCCCGTTCCAAGATCTACATCGAGATGGCCTGCGGAGACCACCTCGTCGTCAAC<br>AACTCCCGTTCCTCCCGTACCGCCCGTGCCCTCCGTCACCACAAGTACCGTAAGACCTGCAAGCGTTGC<br>CGTGTCTCCGACGAGGACCTCAACAAGTTCCTCACCAAGGCCAACGAGGACCAAACCTCCGTCAAGGT<br>CAAGGTCGTCTCCGCCCCAACCCGTACCAAGAAGGTAAGTTTAAACATATcTATACTAACTAACCCTGA<br>TTATTTAAATTTTCAGGCCATGCCAAAGTCCGTCGCCCGTGCCCCAAAGCCACTCGAGAACACCGAGG<br>CCGCCCAAGCCCAACCATCCGGATCCAAGTTCTCCCCAGCCATCCCAGTCTCCACCCAAGAGTCCGTCT<br>CCGTCCCAGCCTCCGTCTCCACCTCCATCTCCTCCATCTCCACCGGAGCCACCGCCTCCGCCCTCGTCAA<br>GGGAAACACCAACCCAATCACCTCCATGTCCGCCCCAGTCCAAGCCTCCGCCCCAGCCCTCACCAAGTC<br>CCAAACCGACCGTCTCGAGGTCCTCCTCAACCCAAAGGACGAGATCTCCCTCAACTCCGGAAAGCCAT<br>TCCGTGAGCTCGAGTCCGAGCTCCTCTCCCGTCGTAAGGTAAGTTTAAACAGTTCGGTACTAACTAACC<br>ATACATATTTAAATTTTCAGAAGGACCTCCAACAAATCTACGCCGAGGAGCGTGAGAACTACCTCGGA<br>AAGCTCGAGCGTGAGATCACCCGTTTCTTCGTCGACCGTGGATTCCTCGAGATCAAGTCCCCAATCCTC<br>ATCCCACTCGAGTACATCGAGCGTATGGGAATCGACAACGACACCGAGCTCTCCAAGCAAATCTTCCG<br>TGTCGACAAGAACTTCTGCCTCCGTCCAATGCTCGCCCCAAACttcTACAACTACatgCGTAAGCTCGACC<br>GTGCCCTCCCAGACCCAATCAAGATCTTCGAGATCGGACCATGCTACCGTAAGGAGTCCGACGGAAA<br>GGTAAGTTTAAACATGATTTTACTAACTAACTAATCTGATTTAAATTTTCAGGAGCACCTCGAGGAGTT<br>CACCATGCTCggaTTCggaCAAATGGGATCCGGATGCACCCGTGAGAACCTCGAGTCCATCATCACCGA<br>CTTCCTCAACCACCTCGGAATCGACTTCAAGATCGTCGGAGACTCCTGCATGGTCttcGGAGACACCCT<br>CGACGTCATGCACGGAGACCTCGAGCTCTCCTCCGCCGTCGTCGGACCAATCCCACTCGACCGTGAGT<br>GGGGAATCGACAAGCCATGGATCGGAGCCGGATTCGGACTCGAGCGTCTCCTCAAGGTCAAGCACGA<br>CTTCAAGAACATCAAGCGTGCCGCCCGTTCCGAGTCCTACTACAACGGAATCTCCACCAACCTCTAAAC<br>CCAGCTTTCTTGTACAAAGTGGTCCCC |

### Supplementary Table 3 – Transgenic *C. elegans* Strains

| Name | Description | Genetic background | Plasmid 1 | Plasmid 2 | Plasmid 3 |
|---|---|---|---|---|---|
| SGR57 | Psur-5::HuPKIαNES::MmNPYRS; Prpr-1::tRNA(C15); Prps-0::GFP(am)::mCherry::HA::EGL-13NLS | smg-6 | JOS97 | ZX297 | SG88 |
| SGR58 | Psur-5::HuPKIαNES::MmNPYRS; Prpr-1::tRNA(C15); Psur-5::enhancer(Y37am)::mCherry::SL2::GFP::EGL-13NLS | N2 | JOS97 | ZX297 | JOS94 |
| SGR59 | Psur-5::HuPKIαNES::MmNPYRS; Prpr-1::tRNA(C15); Psur-5::mCherry::SL2::GFP::EGL-13NLS | N2 | JOS97 | ZX297 | JOS77 |
| SGR60 | Psur-5::HuPKIαNES::MmNPYRS; Prpr-1::tRNA(C15); Psur-5::enhancer::mCherry::SL2::GFP::EGL-13NLS | N2 | JOS97 | ZX297 | JOS52 |
| SGR61 | Psur-5::HuPKIαNES::MmNPYRS; Prpr-1::tRNA(C15); Psur-5::enhancer::mCherry::SL2::EGL-13NLS::BFP | N2 | JOS97 | ZX297 | JOS233 |
| SGR62 | Psur-5::HuPKIαNES::MmNPYRS; Prpr-1::tRNA(C15); Psur-5::enhancer(Y37am, R35A)::mCherry::SL2::GFP::EGL-13NLS | N2 | JOS97 | ZX297 | JOS137 |
| SGR63 | Psur-5::HuPKIαNES::MmNPYRS; Prpr-1::tRNA(C15); Psur-5::enhancer(Y37am, E103A)::mCherry::SL2::GFP::EGL-13NLS | N2 | JOS97 | ZX297 | JOS145 |
| SGR64 | Psur-5::HuPKIαNES::MmNPYRS; Prpr-1::tRNA(C15); Psur-5::enhancer(Y37am, R35A, E103A)::mCherry::SL2::GFP::EGL-13NLS | N2 | JOS97 | ZX297 | JOS140 |
| SGR65 | Psur-5::HuPKIαNES::MmNPYRS; Prpr-1::tRNA(C15); Psur-5::enhancer(Y37am, W47A)::mCherry::SL2::GFP::EGL-13NLS | N2 | JOS97 | ZX297 | JOS142 |
| SGR66 | Psur-5::HuPKIαNES::MmNPYRS; Prpr-1::tRNA(C15); Psur-5::minimiser(Y116am)::mCherry::HA::SL2::GFP::EGL-13NLS | smg-6 | JOS97 | SE150 | JOS229 |
| SGR67 | Psur-5::HuPKIαNES::MmNPYRS; Prpr-1::tRNA(C15); Psur-5::minimiser(Y116am, D98A)::mCherry::HA::SL2::GFP::EGL-13NLS | smg-6 | JOS97 | SE150 | JOS230 |
| SGR68 | Psur-5::HuPKIαNES::MmNPYRS; Prpr-1::tRNA(C15); Psur-5::minimiser(Y116am, D119A)::mCherry::HA::SL2::GFP::EGL-13NLS | smg-6 | JOS97 | SE150 | JOS232 |
